# Development of an Exploratory Taxonomy of Veterinary AI Query Types and Stage-Specific Query Preferences: An Expert Panel Study

**DOI:** 10.64898/2026.06.07.730654

**Authors:** Chan Huh, Hyeyeon Huh, Jinsu Ahn, Minchul Park

**Author notes:** These authors contributed equally to this work (Chan Huh, Hyeyeon Huh).

## Abstract

**Background/Objectives:** The integration of large language model (LLM)-based AI tools into veterinary clinical practice is rapidly increasing; however, veterinary-specific frameworks for organizing AI query types and examining stage-specific preferences remain limited. This study aimed to develop an exploratory taxonomy of veterinary AI query types and to characterize veterinarians’ stage-specific query preferences through a structured expert-panel survey.

**Methods:** Candidate query types were drafted with the assistance of a locally deployed LLM from a preliminary review of de-identified veterinary AI chatbot query logs and refined through literature review. In a structured online survey, 43 veterinarians then selected the Clinical Support query types (A–H) they would prefer to direct to AI at each of five clinical stages and indicated preferred response formats; the survey measured self-reported query preferences, not observed use.

**Results:** A taxonomy of 3 categories and 21 subtypes was established: Clinical Support Queries (Types A–H), Evidence-Based Research Queries (Types I–L), and Terminology and Drug Reference Queries (Types M–U). Across stages, Type B (Differential Reasoning) was most often ranked first (64/213 first-choice responses), and Type D (Clinical Decision Support) was most often ranked first for the post-consultation stage (21/42, 50.0%). In analyses accounting for repeated measures using generalized estimating equations, respondents with ≥10 years of experience more often ranked Type G (Evidence Search) first than those with <10 years (OR 3.09, 95% CI 1.11–8.59; unadjusted p = 0.031), but this contrast was not significant after Benjamini–Hochberg adjustment across four contrasts (adjusted p = 0.062). Respondents from university/teaching hospitals (n = 7) ranked Type G first in 14 of 33 responses, compared with 10 of 135 responses from secondary referral hospitals (OR 9.27, 95% CI 2.94–29.22; adjusted p < 0.001), although this estimate is imprecise because the subgroup is small.

**Conclusions:** This study proposes a veterinary-specific exploratory taxonomy of AI query types and characterizes how respondents’ first-choice preferences varied across clinical stages. These findings offer a preliminary, exploratory framework that may inform the future design of context-aware, stage-adaptive veterinary AI systems, pending cross-national replication and formal validation.

## Introduction

The integration of large language model (LLM)-based AI tools into veterinary clinical practice is rapidly increasing. LLMs are AI systems trained on large text corpora to generate human-like responses to natural-language questions, and are typically accessed through conversational chatbot interfaces. Veterinarians are increasingly using generative AI tools such as ChatGPT and Claude for clinical assistance, documentation, and academic information retrieval (Chu, 2024). A 2024 survey of veterinary professionals across the United States and Canada found that approximately 70% reported using AI tools at least weekly (Gabor et al., 2025).

In human medicine, studies on physician LLM usage patterns have begun to accumulate. A systematic review of 519 healthcare LLM evaluation studies found that clinical reasoning support, differential diagnosis, and documentation were among the dominant AI application categories in medical practice (Bedi et al., 2025). A large-scale quantitative study of 989 physicians analyzing 106,942 query-and-answer pairs found that clinical support and research were the dominant use cases, with significant variation across demographic groups (Qiu et al., 2025). A global survey of healthcare professionals similarly reported widespread use of AI for evidence-based guideline access, diagnostic suggestions, and documentation (Ozkan et al., 2025).

However, these studies share a critical limitation: they do not distinguish query types by clinical stage (pre-, intra-, and post-consultation), failing to capture the context-dependent nature of clinical information needs. Furthermore, they target human medicine and do not reflect the unique characteristics of veterinary practice, including species- and breed-specific disease patterns, the absence of patient self-report, and distinct diagnostic and treatment protocols.

Research on physician clinical information-seeking behavior has established that information needs differ systematically by clinical stage and experience level (Daei et al., 2020), and that experienced physicians have more complex, evidence-gap-related information needs compared to less experienced colleagues (Ostropolets et al., 2020). These principles suggest that a multidimensional taxonomy reflecting clinical context may support the design of veterinary AI systems.

Existing veterinary AI research has focused primarily on LLM performance on standardized examinations (Alonso Sousa et al., 2025) and general adoption surveys (Gabor et al., 2025), with, to our knowledge, no prior study systematically classifying the types of queries veterinarians direct to AI systems in clinical practice. Such a taxonomy may be valuable for designing clinically appropriate AI systems and establishing rigorous benchmark evaluation frameworks.

An exploratory taxonomy was first developed from a preliminary review of de-identified veterinary chatbot queries and refinement against the literature. The resulting taxonomy was subsequently used in a structured survey of 43 veterinarians to characterize stage-specific query preferences and response requirements. The analysis describes self-reported preferences across clinical stages, experience levels, and institution types; it does not describe observed query behavior, the reasons for querying, or clinical effects.

## Materials and Methods

### Study Design

This exploratory three-phase study combined a preliminary qualitative review of de-identified chatbot queries, literature-based refinement of the resulting candidate types into a taxonomy, and a cross-sectional expert-panel survey. Two separately collected data sources served different purposes: pre-existing de-identified chatbot query logs informed the candidate survey items (Phase 1), and the expert-panel survey provided the participant characteristics and all stage-specific preference data analyzed in this study (Phase 3). Survey responses were not linked to individual chatbot logs, and no clinical-stage or experience subgroup was inferred from the logs. The study protocol was approved through expedited review by the Institutional Review Board of Changwon National University, Republic of Korea, on 11 September 2026 (approval no. 7001066-202607-HR-053). The approved protocol described the previously completed questionnaire-development phases (Phases 1 and 2) and the subsequent anonymous survey. All 43 responses included in the analysis were collected after approval, with data collection completed on 19 September 2026. Under the approved consent procedure, the full participant information sheet was presented on the first survey screen, and voluntary completion and submission constituted consent to participate. No separate signed consent form or consent checkbox was used.

### Information obtained from each data source

The chatbot logs contributed only the de-identified text of submitted queries, which was used to draft candidate query types (Phase 1) and, after the taxonomy was finalized, to check that each subtype was represented among real queries; no participant characteristics and no frequency estimates were taken from them. The survey contributed all participant characteristics (institution type, clinical experience, and clinical specialty, each self-reported) and all outcome data. Because the survey was anonymous and was not linked to chatbot accounts, it cannot be determined whether any survey respondent had also used the chatbot.

### Participants

Forty-three veterinarians from South Korea participated in the anonymous online survey. The protocol targeted approximately 40 participants, with an allowable range of 30–60, to support exploratory characterization of stage-specific preferences while seeking diversity in institution type, clinical experience, and specialty. Recruitment used a publicly posted invitation in a veterinarian-operated online chat community, with the moderator’s permission. Participation was voluntary, and researchers did not individually approach potential participants. Eligibility required a veterinary license and active clinical practice at the time of the survey; these were assessed by self-report at the start of the questionnaire, and respondents answering otherwise were routed out before the main items. All analyzed respondents, including the two affiliated with research institutions, confirmed current clinical practice at that screen. The investigators and Choonok Company employees were excluded. Participants were asked to respond once; login-based restrictions and IP collection were not used to preserve anonymity. Institution type, clinical experience, and clinical specialty were self-reported. Participant characteristics were as follows. By institution type: secondary referral hospitals (n = 27, 62.8%), university/teaching hospitals (n = 7, 16.3%), primary care clinics (n = 7, 16.3%), and research institutions (n = 2, 4.7%). Mean clinical experience was 8.9 years (median 10.0, range 0.5–20 years); 20 participants (46.5%) had less than 10 years of experience and 23 (53.5%) had 10 or more years. Selected self-reported clinical specialties (multiple responses permitted; related free-text responses grouped) were: internal medicine and related subspecialties (n = 26), surgery (n = 9), diagnostic imaging (n = 6), and emergency/critical care (n = 5). The rules used to convert free-text experience entries to years and to group the specialty responses are given in Supplementary Material S3.

### Taxonomy Development Process

The taxonomy was developed through a three-phase process.

#### Phase 1 — Preliminary Draft Generation from Query Logs

To generate an initial pool of candidate query types for the survey instrument, real-world queries submitted by veterinarians to a RAG-based veterinary clinical AI chatbot were reviewed. The chatbot was built on the ChatGPT API (OpenAI) with a custom retrieval-augmented generation (RAG) pipeline and proprietary search engine developed by Choonok Company, and was deployed as a proof-of-concept system at three animal hospitals in Ulsan and Gyeongsangnam-do, in the south-east of the Republic of Korea (one secondary referral hospital, one cancer center, and one primary-care clinic). Veterinarians at these sites entered free-text clinical questions in a web-based chat interface and received answers generated from passages retrieved from the system’s reference sources, with the retrieved sources listed alongside the answer; a schematic of that interface is given in Supplementary Material S6. Account registration details were not extracted or analyzed for this study; all participant characteristics reported here come from the survey. Because these logs originated from a limited set of proof-of-concept deployment sites, they were used only as a preliminary source for drafting candidate query categories, and not as a basis for any frequency, prevalence, or usage claims.

Under fixed extraction conditions, 1,785 queries submitted from 34 user accounts in 972 conversations between 1 August 2025 and 30 March 2026 were retrieved from the operating database; the count comprises messages written by users, not chatbot replies, and excludes internal accounts and user-deleted conversations. Accounts did not correspond one-to-one to individuals: a shared account was used at one site so that clinicians could query the system without registering personally, and that account contributed 776 of the 1,785 queries (43.5%). The number of distinct veterinarians represented is therefore unknown.

All queries were de-identified by a Choonok Company developer who is not an author of this study. The de-identification used a locally deployed large language model (nvidia/Gemma-4-26B-A4B-NVFP4, temperature 0) served with vLLM on a local server, which masked direct identifiers (personal names, contact numbers, and chart or account numbers) and quasi-identifiers (hospital and institution names, place names, and patient-animal names) while preserving drug, disease, test, breed, and species names as analytic content. Extracted identifiers were verified against the source text, re-checked in a separate adjudication step, and screened by an exhaustive scan against a list of 37 known real names. All de-identification processing was performed locally and no data were transmitted externally. Only the de-identified corpus was used in this study, and no account-identifying information was used at any stage. The deployed system was a non-commercial proof-of-concept developed to support clinical care, and its users had agreed, under the system’s privacy policy, to the use of their data for service improvement and research; all queries were additionally de-identified before any analysis.

The de-identified queries were then used as a preliminary qualitative source for drafting candidate query types with the assistance of the locally deployed model described above. The research team reviewed the generated candidates for duplication, clinical relevance, and scope, merging, splitting, renaming, or removing preliminary types as needed. The resulting candidate set was then taken forward to Phase 2 for literature-based refinement. In this manuscript the terms are used deliberately and not interchangeably: “AI chatbot” denotes the deployed RAG-based system, “large language model (LLM)” an underlying generative model, “RAG” the retrieval-augmented-generation pipeline, and “AI query” an information request submitted by a professional.

#### Phase 2 — Literature-Based Taxonomy Refinement

The candidate query types generated in Phase 1 were reviewed against the relevant literature and refined by the research team into an exploratory taxonomy comprising three categories and 21 subtypes: Clinical Support Queries (Types A–H), Evidence-Based Research Queries (Types I–L), and Terminology and Drug Reference Queries (Types M–U). The three categories are distinguished by the primary intent of the query rather than by clinical discipline, and the definition and an illustrative example of each of the 21 subtypes are given in Table 2 of the Results; the definitions as supplied to the classifier are reproduced in Supplementary Material S2. All four authors independently reviewed the proposed subtype definitions and category assignments, and disagreements were resolved through discussion before the taxonomy was finalized for the survey.

**Table 1.**
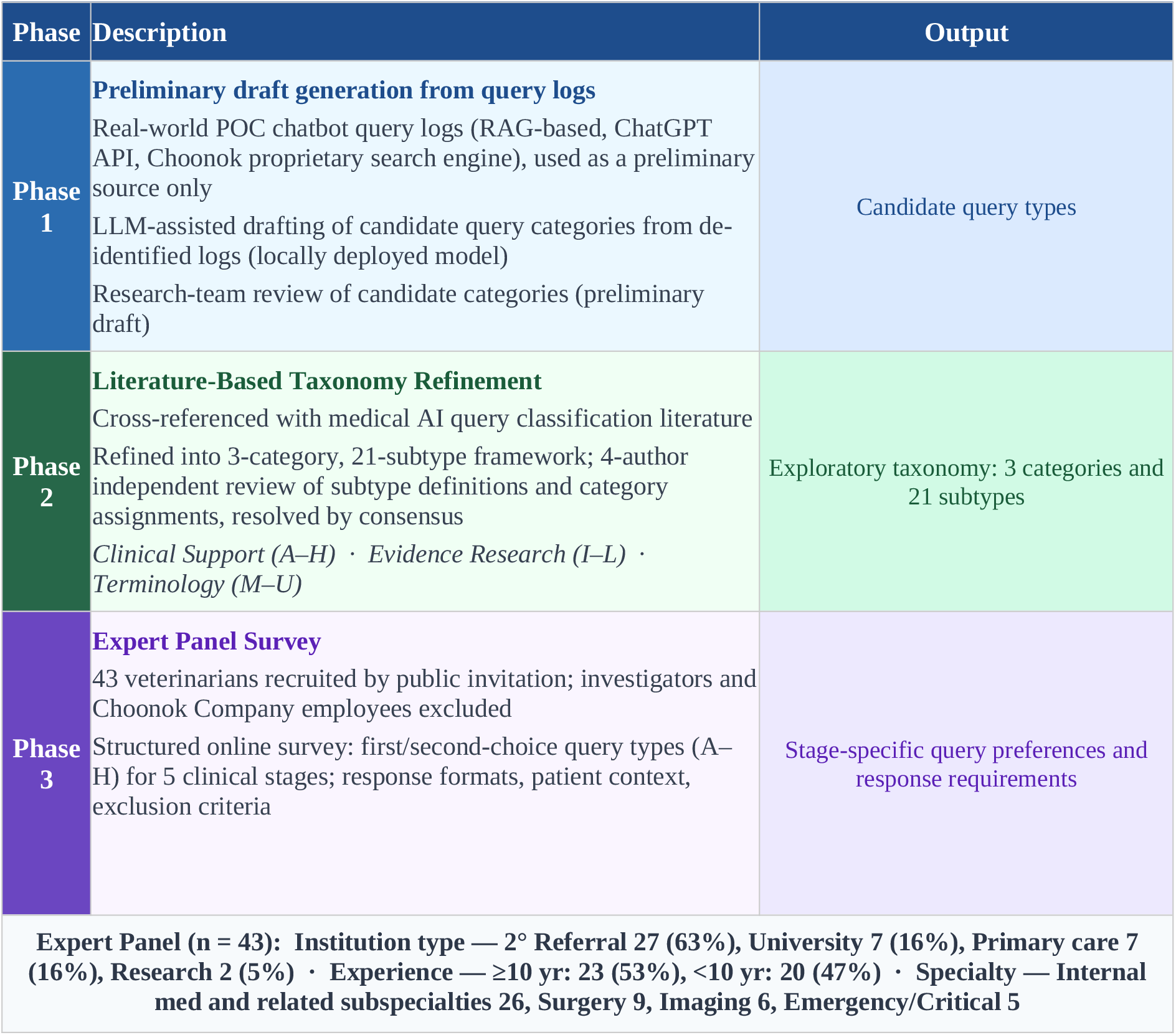
Three-Phase Taxonomy Development Process.

| Phase | Description | Output |
|---|---|---|
| **Phase 1** | **Preliminary draft generation from query logs**<br>Real-world POC chatbot query logs (RAG-based, ChatGPT API, Choonok proprietary search engine), used as a preliminary source only<br>LLM-assisted drafting of candidate query categories from de-identified logs (locally deployed model)<br>Research-team review of candidate categories (preliminary draft) | Candidate query types |
| **Phase 2** | **Literature-Based Taxonomy Refinement**<br>Cross-referenced with medical AI query classification literature<br>Refined into 3-category, 21-subtype framework; 4-author independent review of subtype definitions and category assignments, resolved by consensus<br>*Clinical Support (A–H)* · *Evidence Research (I–L)* · *Terminology (M–U)* | Exploratory taxonomy: 3 categories and 21 subtypes |
| **Phase 3** | **Expert Panel Survey**<br>43 veterinarians recruited by public invitation; investigators and Choonok Company employees excluded<br>Structured online survey: first/second-choice query types (A–H) for 5 clinical stages; response formats, patient context, exclusion criteria | Stage-specific query preferences and response requirements |
| **Expert Panel (n = 43): Institution type — 2° Referral 27 (63%), University 7 (16%), Primary care 7 (16%), Research 2 (5%) · Experience — ≥10 yr: 23 (53%), <10 yr: 20 (47%) · Specialty — Internal med and related subspecialties 26, Surgery 9, Imaging 6, Emergency/Critical 5** |  |  |

**Table 2.**
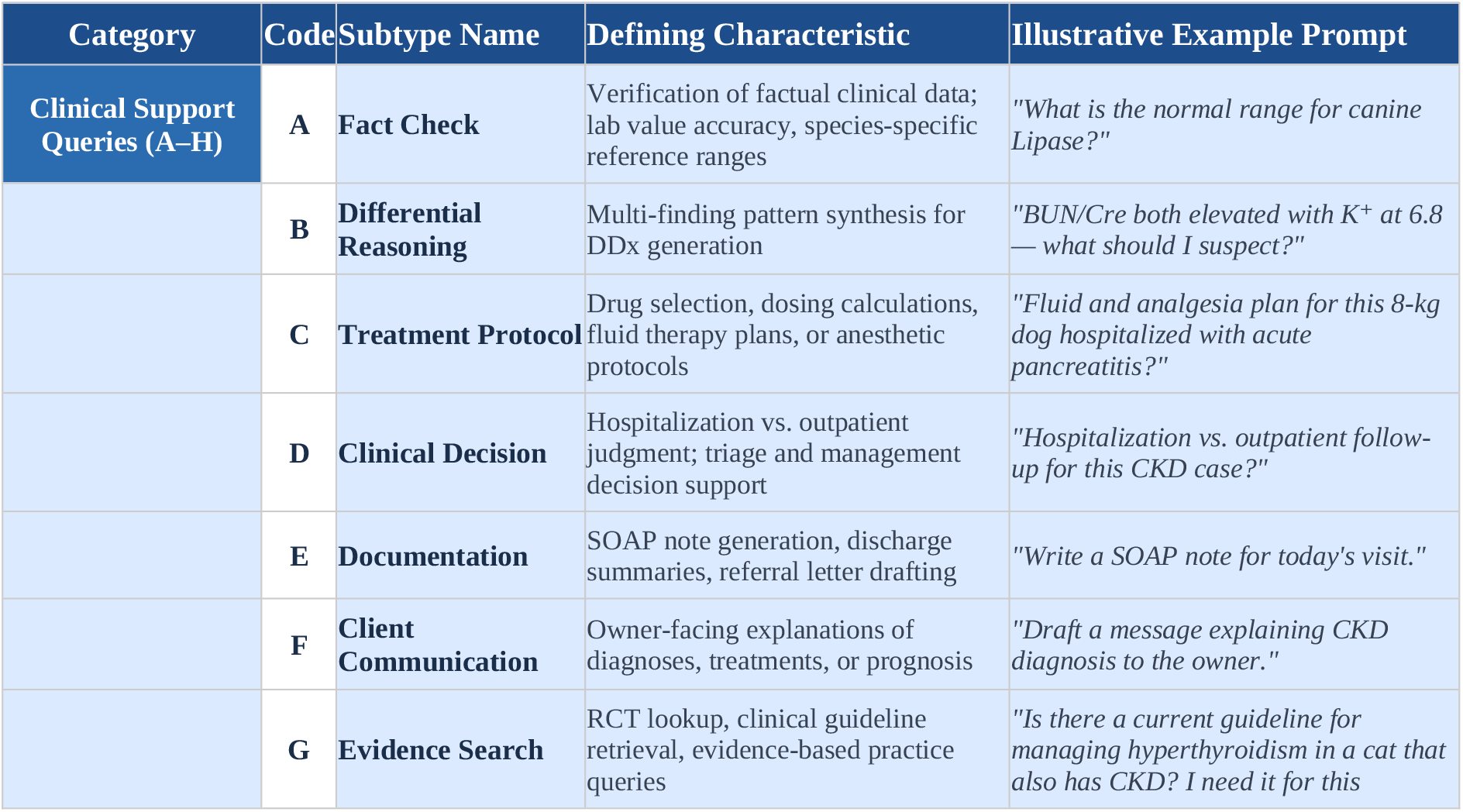
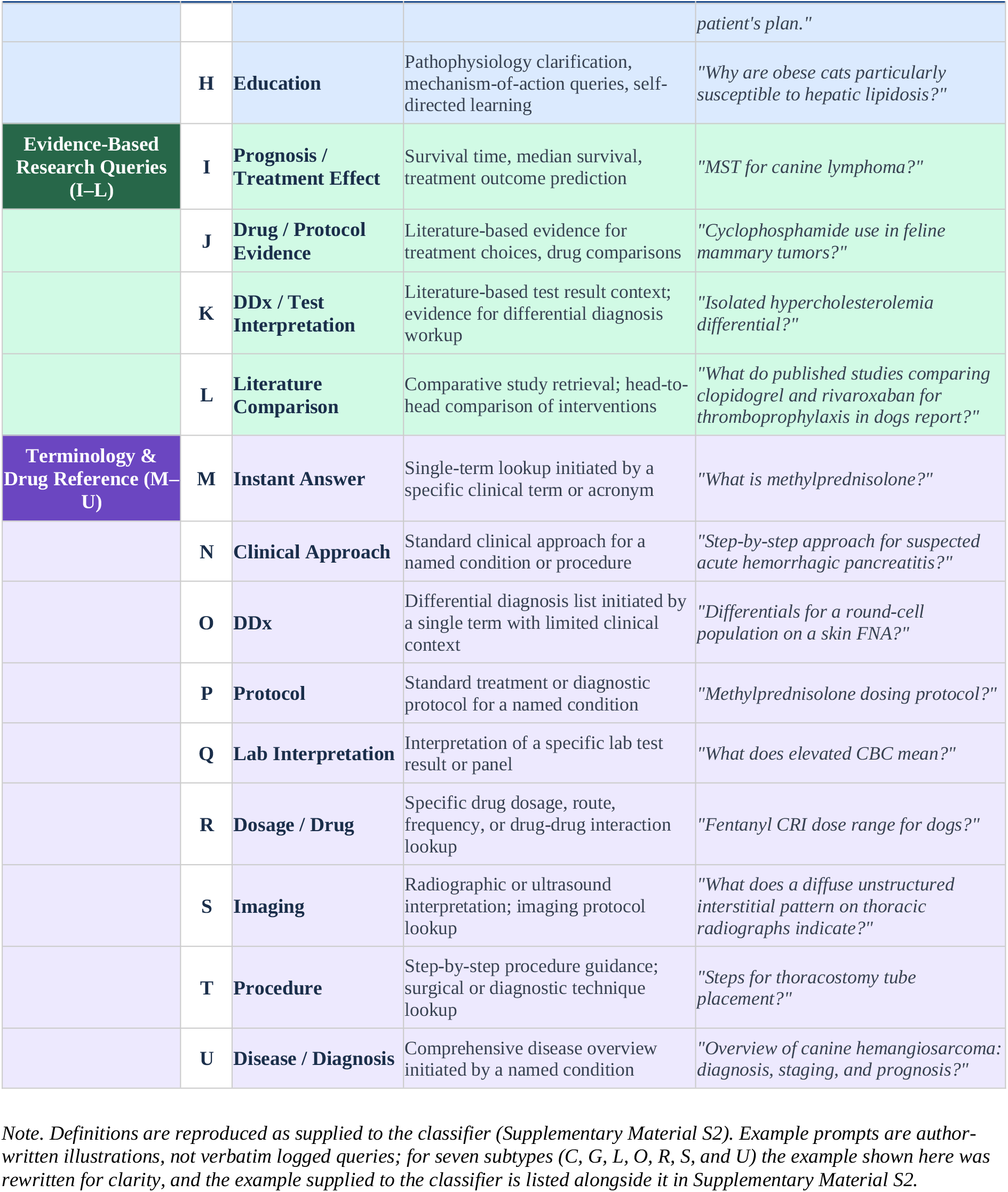
Veterinary AI Query Taxonomy with Illustrative Example Prompts and Operational Definitions (3 Categories, 21 Subtypes)

#### Phase 3 — Expert Panel Survey

The exploratory taxonomy established in Phase 2 was presented to 43 veterinarians through a structured online survey to characterize stage-specific query preferences, preferred response formats, patient-context requirements, and training-data exclusion criteria. Participants selected their first-and second-choice query types among the eight Clinical Support types (A–H) for each of five clinical stages. The five stages follow the course of a clinical encounter — pre-consultation, intra-consultation before diagnostic testing, intra-consultation after testing, post-consultation, and off-duty — and are set out with the full item wording in the Survey Instrument section below. Descriptive analyses summarized query-type rankings and response-format selections. These preference questions did not directly measure agreement with subtype validity. Phase 3 was designed to characterize veterinarians’ preferences rather than to establish consensus on the validity of the taxonomy. Accordingly, no formal consensus procedure such as Delphi or the RAND/UCLA appropriateness method (Murphy et al., 1998; Fitch et al., 2001), and no psychometric validation metric such as inter-rater reliability or a content validity index, was applied to the survey responses; this is consistent with the exploratory scope of the present study (Elo and Kyngäs, 2008). Formal inter-rater reliability metrics and CVI calculation are recommended for future confirmatory studies in which the taxonomy is applied to independent datasets.

#### Coverage check using the exploratory taxonomy

After the taxonomy had been established in Phase 2, it was applied to the same 1,785 de-identified user queries as an internal coverage check, to see whether each subtype was represented among real queries. Because the corpus had contributed to candidate-type generation, this procedure was not treated as independent validation. Classification was run locally with nvidia/Gemma-4-26B-A4B-NVFP4 (temperature 0) in two stages that mirrored the taxonomy’s hierarchy — category, then subtype within that category — each read from the model’s output-token log-probabilities, with a single primary-intent label per query and a separate code for queries containing no clinical information request. Each assignment was verified against the assigned subtype’s definition in a yes/no judgment, with up to five reclassification iterations. After strict verification caused queries to cycle among the closely defined Evidence-Based Research subtypes (I–L), the verification criterion was relaxed to reject an assignment only when the subtype clearly did not fit, accepting assignments that merely competed with an adjacent subtype; the prompt that was retained applies this relaxed criterion to all subtypes, and the earlier, stricter wording was not retained. The prompts, which were written in Korean, are reproduced from the pipeline with English translations in Supplementary Material S1, together with the run configuration and the mapping used to read the category decision; the definitions and examples supplied to the model are given in Supplementary Material S2. Queries that did not converge within five iterations (128 of 1,785; 7.2%) were independently adjudicated by all four authors; only the final labels were retained, so the number of labels the authors changed was not recorded. The remaining 1,657 labels were not reviewed by humans. All 21 subtypes were represented among the classified queries. No frequency, prevalence, or distributional results are derived from this step, and no inter-rater reliability was estimated.

### Survey Instrument

After two eligibility questions and three background items (institution type, clinical experience, and clinical specialty), the survey comprised seven thematic modules. Module 1 (Clinical L1) asked respondents to select their first- and second-choice query types among Types A–H for each of five clinical stages — pre-consultation, intra-consultation (pre-test), intra-consultation (post-test), post-consultation, and off-duty — along with preferred response formats per type. This five-stage framework was developed by the research team based on the established principle that clinical information needs differ systematically across phases of the clinical encounter (Daei et al., 2020), and refined to reflect the specific workflow of veterinary practice, in which an off-duty stage captures AI use for continuing education and case review outside active consultation hours. Module 2 (Clinical L2) assessed essential patient context variables. Module 3 (Clinical L3) assessed training data exclusion criteria. Modules 4–5 (Research L1–L2) assessed evidence-based query preferences and response formats. Module 6 (Research L3) assessed research data quality criteria. Module 7 (Terminology L1) assessed drug and terminology query preferences and preferred response ordering. Stage-specific rankings were collected only for the eight Clinical Support types (A–H), so all stage-level distributions and GEE analyses in this article refer to Types A–H rather than to all 21 subtypes. The main text reports Module 1 and the multiple-selection items of Modules 2 and 3; descriptive results for Modules 4–7 and the second-choice distributions are given in Supplementary Material S4, and free-text items were not coded or analyzed. The questionnaire as deployed, with an English translation, is provided in Supplementary Material S3.

### Structure of the Exploratory Taxonomy

The exploratory taxonomy comprises 3 categories and 21 subtypes, as presented in Table 2. The categorization follows the primary intent of each query: whether the immediate goal is to support a clinical action (Clinical Support Queries, A–H), retrieve or synthesize published evidence (Evidence-Based Research Queries, I–L), or look up a specific term, dosage, or procedure entry point (Terminology and Drug Reference Queries, M–U). In the coverage check, each query was assigned to a single subtype according to its primary intent (point-of-care clinical support, A–H; research and literature appraisal, I–L; or quick terminology/drug reference, M–U), using only the definitions and examples in Supplementary Material S2 and the prompts in Supplementary Material S1. No separate coding manual, list of borderline cases, or tie-break hierarchy was used during classification. Formal inter-rater reliability on an independent sample was not assessed and is identified as a priority for confirmatory work.

### Statistical Analysis

Because each participant responded across all five clinical stages, responses were clustered within participants. To account for this repeated-measures structure, associations between query type and participant characteristics were examined using generalized estimating equations (GEE), with each query type of interest treated as a binary outcome. Three separate binary GEE models with a logit link, an exchangeable working correlation structure, and robust sandwich covariance estimates examined Type G (Evidence Search) and Type D (Clinical Decision Support) by clinical experience (≥10 vs. <10 years), and Type G by institution type (the two research-institute participants were excluded; secondary referral hospitals were the reference). The experience models did not adjust for institution, and the institution model did not adjust for experience.

Types G and D were not pre-specified as outcomes in the approved protocol. Descriptive distributions for all eight Clinical Support types were examined first, and Types G and D were then selected for modelling; the selection was therefore data-driven, and the reported p-values do not account for it. The other six types were not modelled. The permutation tests listed in the protocol were not performed; inference relies on robust GEE Wald tests with Benjamini–Hochberg correction, a deviation from the protocol’s analysis plan. Odds ratios are reported with unadjusted 95% Wald confidence intervals. Benjamini–Hochberg correction was applied across the four reported coefficient contrasts (two experience contrasts and two institution contrasts). A two-degree-of-freedom Wald test additionally evaluated the institution factor as a whole. Sensitivity analyses considered independence working correlation, bias-reduced covariance, adjustment for clinical stage, non-use responses coded as non-selection, and joint adjustment for experience, institution, and clinical stage. A same-sample experience model was fitted to the 41 participants included in the institution analysis. Analyses used statsmodels 0.15.0, and all fitted models converged. Consistent with the exploratory design and the small subgroup sizes, these comparisons are treated as hypothesis-generating rather than confirmatory.

Response accounting was as follows: all 215 stage-level items (43 participants × 5 stages) had a submitted response; 213 first choices mapped to Types A–H. Two responses (one pre-consultation and one post-consultation), both from a single participant with ≥10 years of experience at a university/teaching hospital who selected “other—did not ask,” were excluded from the A–H analyses as non-use responses rather than unanswered items. Analyzable A–H first-choice responses numbered 113 (≥10 years) and 100 (<10 years) by experience, and 135 (secondary referral), 33 (university/teaching), 35 (primary care), and 10 (research institution, excluded from the institution comparison). Stage and subgroup percentages for Types A–H use these denominators; the stage denominators were 42 (pre-consultation), 43 (intra-consultation, pre-test), 43 (intra-consultation, post-test), 42 (post-consultation), and 43 (off-duty). All stage-level analyses concern Types A–H only. Response-format and multiple-selection items use 43 respondents as the denominator; multiple-selection percentages may sum to more than 100%. Primary analyses used first-choice responses, which give a single highest-priority preference per participant at each clinical stage and avoid treating first-and second-choice rankings as equivalent outcomes. Second-choice responses were also incomplete, with 37 to 39 of the 43 participants recording a second choice at each stage. They were therefore retained for descriptive analysis only and are reported in Supplementary Material S4, using the number of valid second-choice responses at each stage as the denominator.

## Results

### Query Type Distribution Across Clinical Stages

First-choice rankings of Types A–H differed descriptively across the five clinical stages (Table 3, Figure 2). Pre-consultation: Type B (Differential Reasoning) was ranked first by 18 participants (42.9%), followed by Type A (21.4%) and Type C (19.0%). Intra-consultation (pre-test): Types B and D were tied as first choice (each 23.3%), followed by Type A (18.6%) and Types C and G (each 16.3%). Intra-consultation (post-test): Type B was first (37.2%), followed by Type C (30.2%). Post-consultation: Type D was the most frequent first choice (21/42, 50.0%). Off-duty: Type B was first (44.2%), followed by Type G (32.6%).

**Figure 1.**
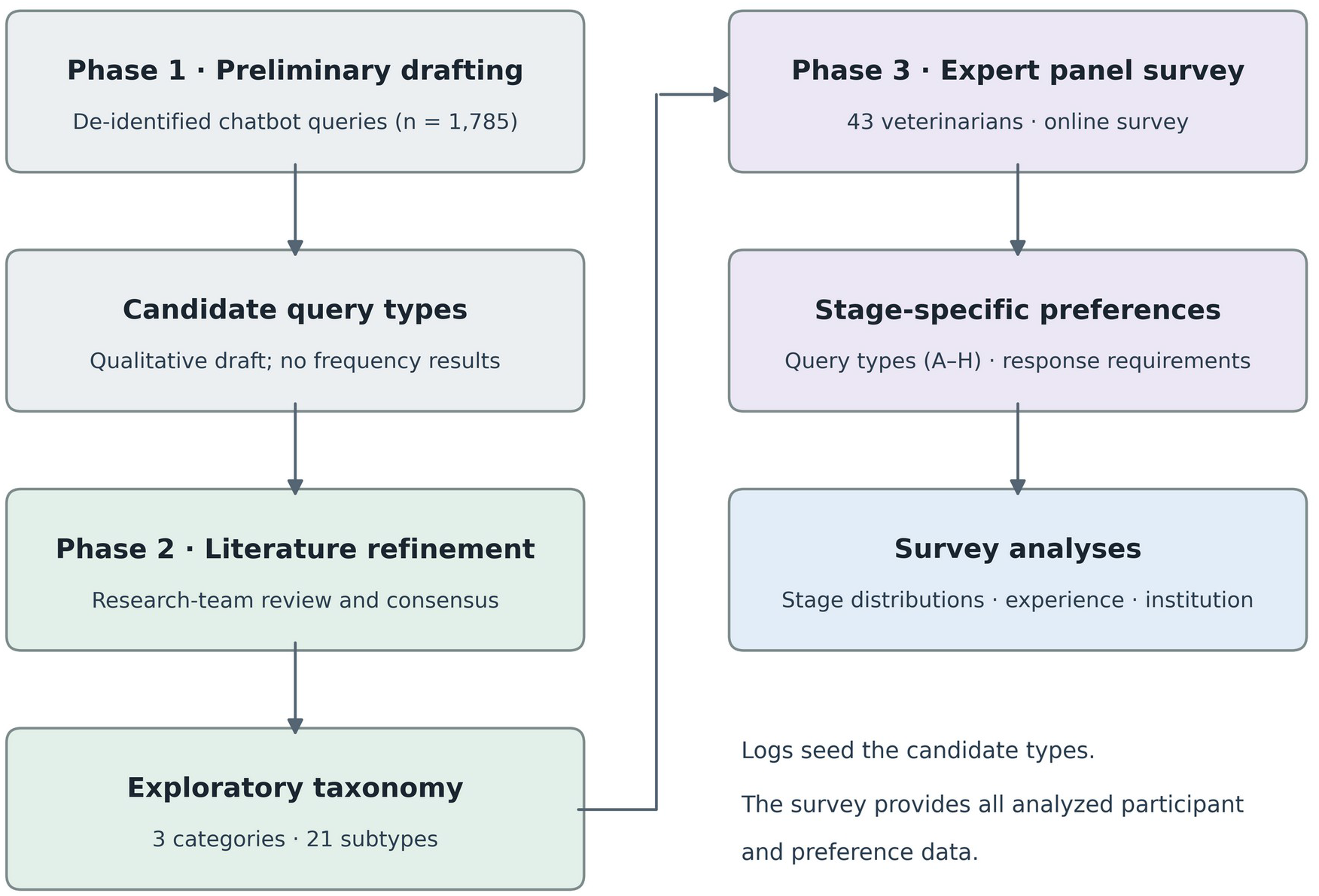
Overview of the study design. Candidate query types were drafted from a preliminary review of de-identified chatbot queries (Phase 1); no frequency or usage results were derived from that phase. The candidates were refined against the literature into the exploratory taxonomy of 3 categories and 21 subtypes (Phase 2), which was then presented to 43 veterinarians in a structured survey (Phase 3). The survey yielded stage-specific query preferences and response requirements for the Clinical Support types (A–H), analyzed by clinical stage, experience level, and institution type using generalized estimating equations.

**Figure 2.**
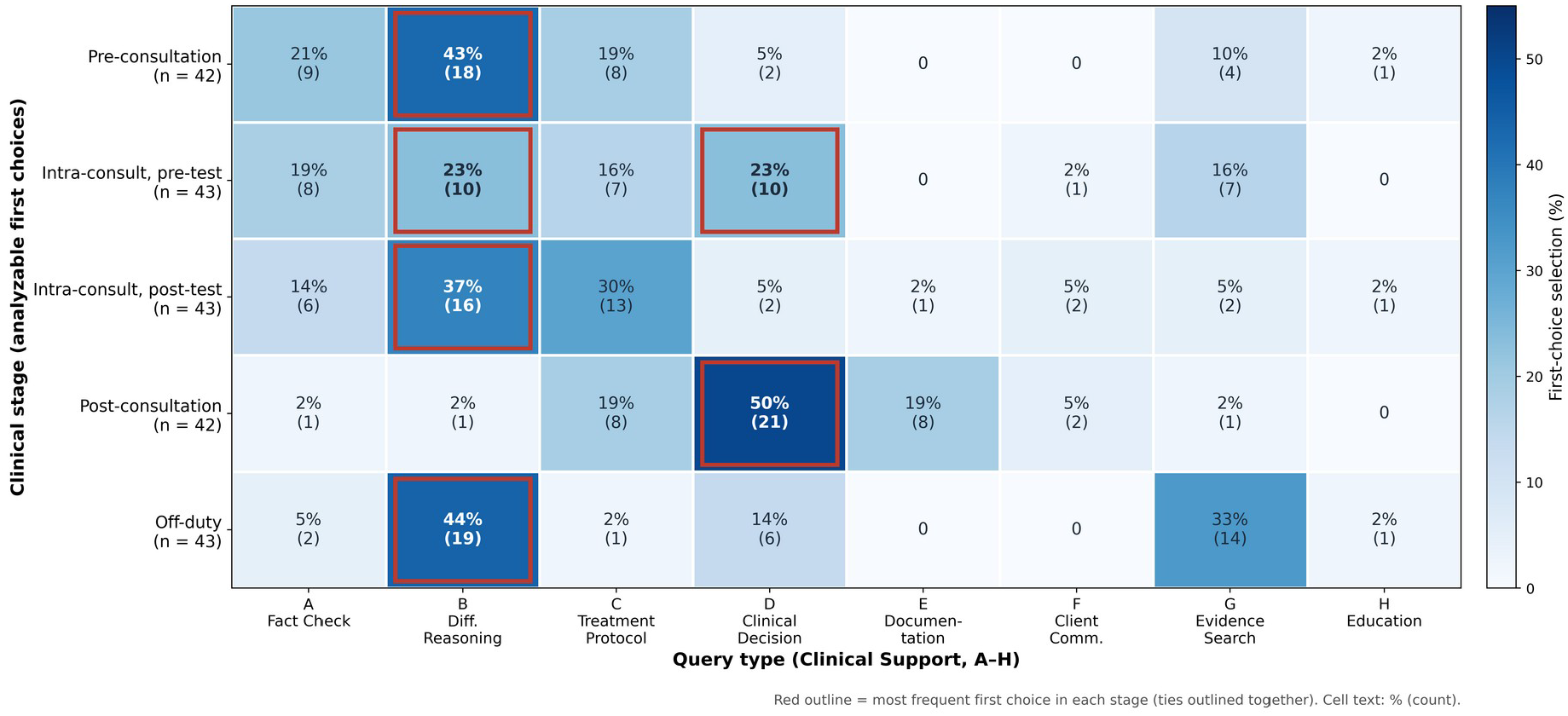
Query Type Distribution Across Clinical Stages. Heat map shows the percentage (and count) of first-choice selections of Clinical Support types A–H by 43 veterinarians across five clinical stages. Percentages use stage-specific denominators of analyzable A–H first choices: 42 (pre-consultation), 43 (intra-consultation, pre-test), 43 (intra-consultation, post-test), 42 (post-consultation), and 43 (off-duty); two “other—did not ask” responses were excluded. Red outlines mark the most frequent first choice in each stage (ties outlined together). Type B (Differential Reasoning) was most often ranked first at pre-consultation, post-test, and off-duty stages, and Type D (Clinical Decision Support) post-consultation (21/42, 50%).

**Table 3.**
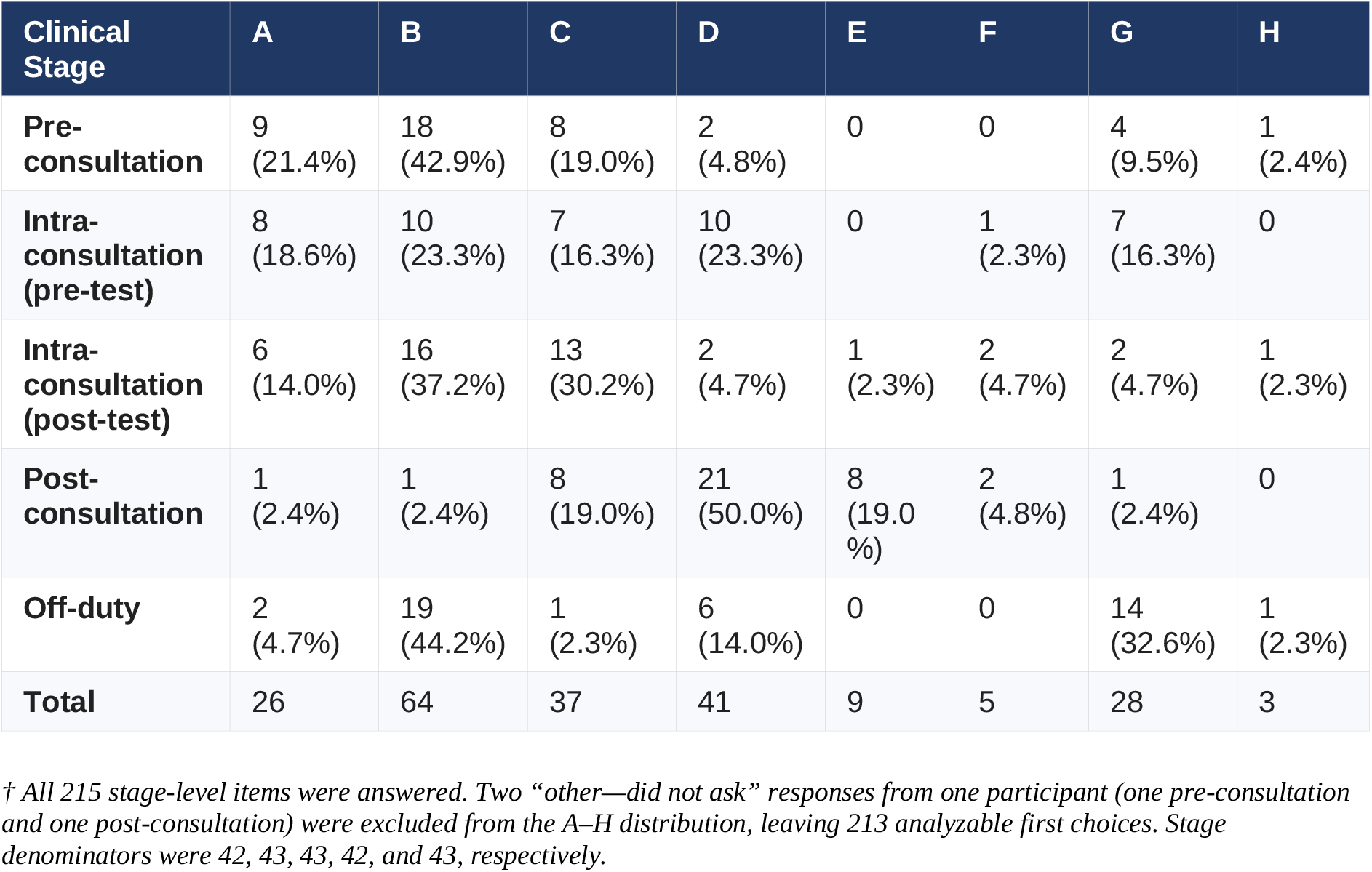
First-Choice Query Type Frequency Matrix by Clinical Stage (43 respondents; stage denominators of analyzable A–H first choices, 42–43)

| **Clinical Stage** | **A** | **B** | **C** | **D** | **E** | **F** | **G** | **H** |
|---|---|---|---|---|---|---|---|---|
| **Pre-consultation** | 9 (21.4%) | 18 (42.9%) | 8 (19.0%) | 2 (4.8%) | 0 | 0 | 4 (9.5%) | 1 (2.4%) |
| **Intra-consultation (pre-test)** | 8 (18.6%) | 10 (23.3%) | 7 (16.3%) | 10 (23.3%) | 0 | 1 (2.3%) | 7 (16.3%) | 0 |
| **Intra-consultation (post-test)** | 6 (14.0%) | 16 (37.2%) | 13 (30.2%) | 2 (4.7%) | 1 (2.3%) | 2 (4.7%) | 2 (4.7%) | 1 (2.3%) |
| **Post-consultation** | 1 (2.4%) | 1 (2.4%) | 8 (19.0%) | 21 (50.0%) | 8 (19.0%) | 2 (4.8%) | 1 (2.4%) | 0 |
| **Off-duty** | 2 (4.7%) | 19 (44.2%) | 1 (2.3%) | 6 (14.0%) | 0 | 0 | 14 (32.6%) | 1 (2.3%) |
| **Total** | 26 | 64 | 37 | 41 | 9 | 5 | 28 | 3 |

*† All 215 stage-level items were answered. Two “other—did not ask” responses from one participant (one pre-consultation and one post-consultation) were excluded from the A–H distribution, leaving 213 analyzable first choices. Stage denominators were 42, 43, 43, 42, and 43, respectively.*

Across all five stages combined (213 analyzable first choices), Type B was most often ranked first (64 responses), followed by D (41), C (37), G (28), and A (26). Second-choice distributions, based on the 37 to 39 participants who recorded a second choice at each stage, are provided in Supplementary Material S4.

### Differences by Clinical Experience

Participants with ≥10 years of experience selected Type G more often than those with <10 years (21/113, 18.6% vs. 7/100, 7.0%; Table 4). In a participant-clustered GEE model, the odds ratio was 3.09 (95% CI 1.11–8.59; unadjusted p = 0.031), with a working correlation of α = 0.138. This association did not meet the 0.05 threshold after Benjamini–Hochberg correction across four reported contrasts (adjusted p = 0.062). Type D selection was numerically more frequent in the less experienced group (22/100, 22.0% vs. 19/113, 16.8%), but the ≥10-year versus <10-year contrast was imprecise and not statistically significant (OR 0.71, 95% CI 0.33–1.56; p = 0.397; adjusted p = 0.530). Both groups showed comparable descriptive frequencies for Types B and C.

**Table 4.** First-Choice Query Type Frequency by Clinical Experience (Q1–Q5 Combined)

| **Query Type** | **≥10 Years (n = 23, responses = 113)** | **<10 Years (n = 20, responses = 100) †** |
|---|---|---|
| **A Fact Check** | 11 (9.7%) | 15 (15.0%) |
| **B Differential Reasoning** | 32 (28.3%) | 32 (32.0%) |
| **C Treatment Protocol** | 19 (16.8%) | 18 (18.0%) |
| **D Clinical Decision Support** | 19 (16.8%) | 22 (22.0%) |
| **G Evidence Search** | 21 (18.6%) | 7 (7.0%) |
| **Other (E, F, H)** | 11 (9.7%) | 6 (6.0%) |

*† Percentages are based on each participant’s five stage-level first-choice responses. Two non-use responses (one pre-consultation, one post-consultation), both from a single ≥10-year participant affiliated with a university/teaching hospital who selected “other (did not ask),” were excluded from the A–H analysis; these were submitted answers, not unanswered items. The ≥10-year group therefore contributed 113 responses and the <10-year group 100 (20 × 5, complete), giving 213 analyzable A–H first-choice responses overall.*

### Differences by Institution Type

Type G accounted for 14 of 33 first-choice responses in university/teaching hospitals (42.4%), compared with 10 of 135 in secondary referral hospitals (7.4%) and 2 of 35 in primary care clinics (5.7%; Table 5, Figure 4). The institution factor was associated with Type G selection in a participant-clustered GEE Wald test (χZ(2) = 16.80, p < 0.001). Relative to secondary referral hospitals, university/teaching affiliation was associated with higher odds of Type G selection (OR 9.27, 95% CI 2.94–29.22; p < 0.001; BH-adjusted p < 0.001), whereas the primary-care contrast was imprecise and not statistically significant (OR 0.76, 95% CI 0.18–3.12; p = 0.701; BH-adjusted p = 0.701). These findings describe an association concentrated in the university/teaching group, not a graded trend across institutions. Estimates remain preliminary given the small university/teaching subgroup (n = 7). The interval was not narrowed under a more conservative variance estimator: with bias-reduced covariance the same contrast gave OR 9.27 (95% CI 2.54–33.87). In a same-sample analysis restricted to the 41 participants in the institution comparison, the experience-only Type G estimate was OR 2.74 (95% CI 0.96–7.86; p = 0.060). In an exploratory model jointly adjusting for experience, institution, and clinical stage, the experience estimate was OR 1.74 (95% CI 0.56–5.39; p = 0.341), while the university/teaching estimate was OR 12.09 (95% CI 2.77–52.69; p < 0.001). These sensitivity p-values are unadjusted; the joint model, with 26 Type G selections and eight regression coefficients including the intercept, has limited information and should not be interpreted as confirmatory.

**Figure 3.**
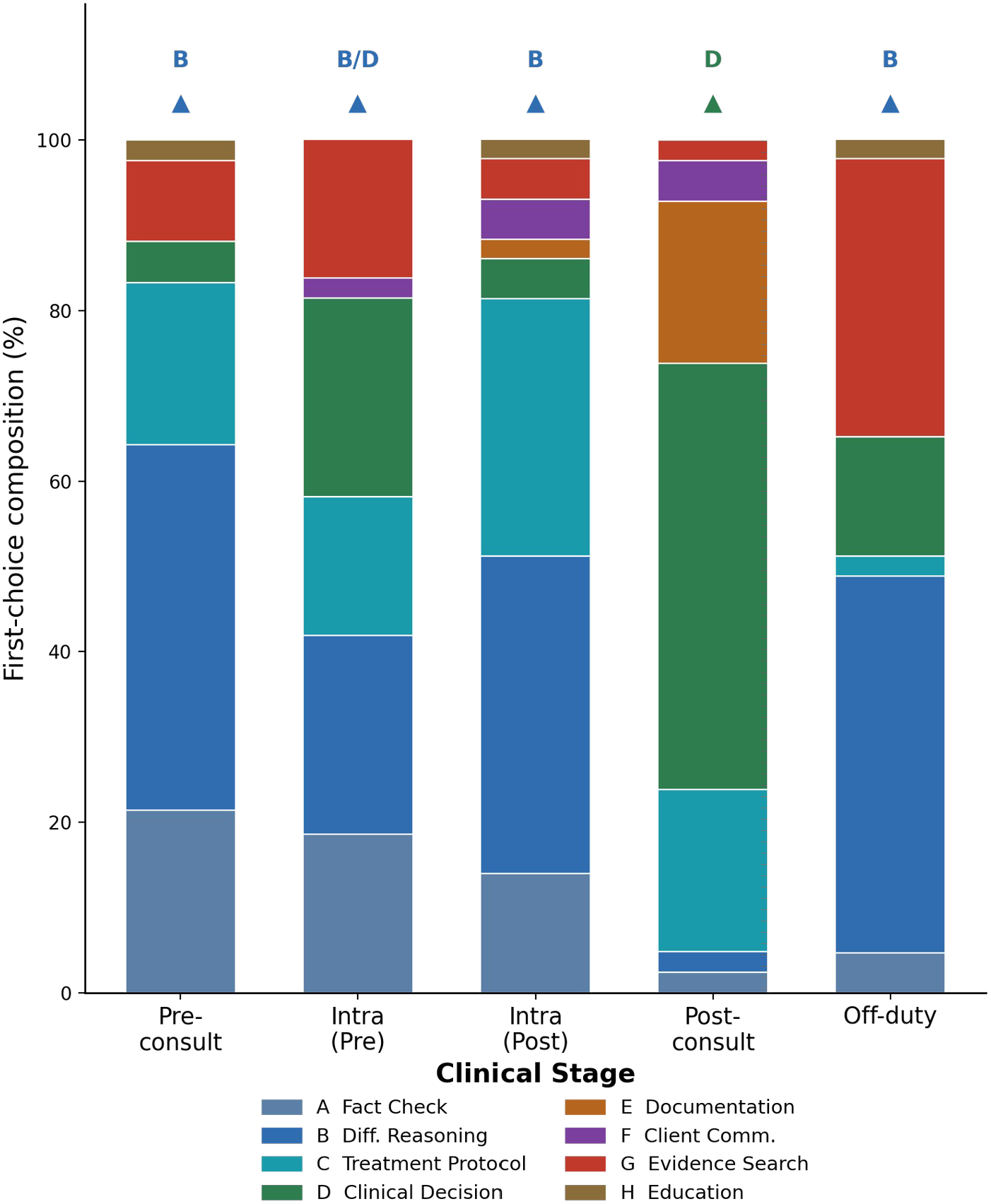
Query Type Composition Across Clinical Stages. Stacked bars represent the proportional distribution of first-choice query type selections across five veterinary clinical stages (43 respondents; stage denominators of analyzable A–H first choices were 42, 43, 43, 42, and 43, respectively). The most frequently selected query type per stage is indicated above each bar (▴). Color coding corresponds to query types consistent with Figure 2. All eight query types (A–H) are displayed; minor types with low frequency appear as narrow bands.

**Figure 4.**
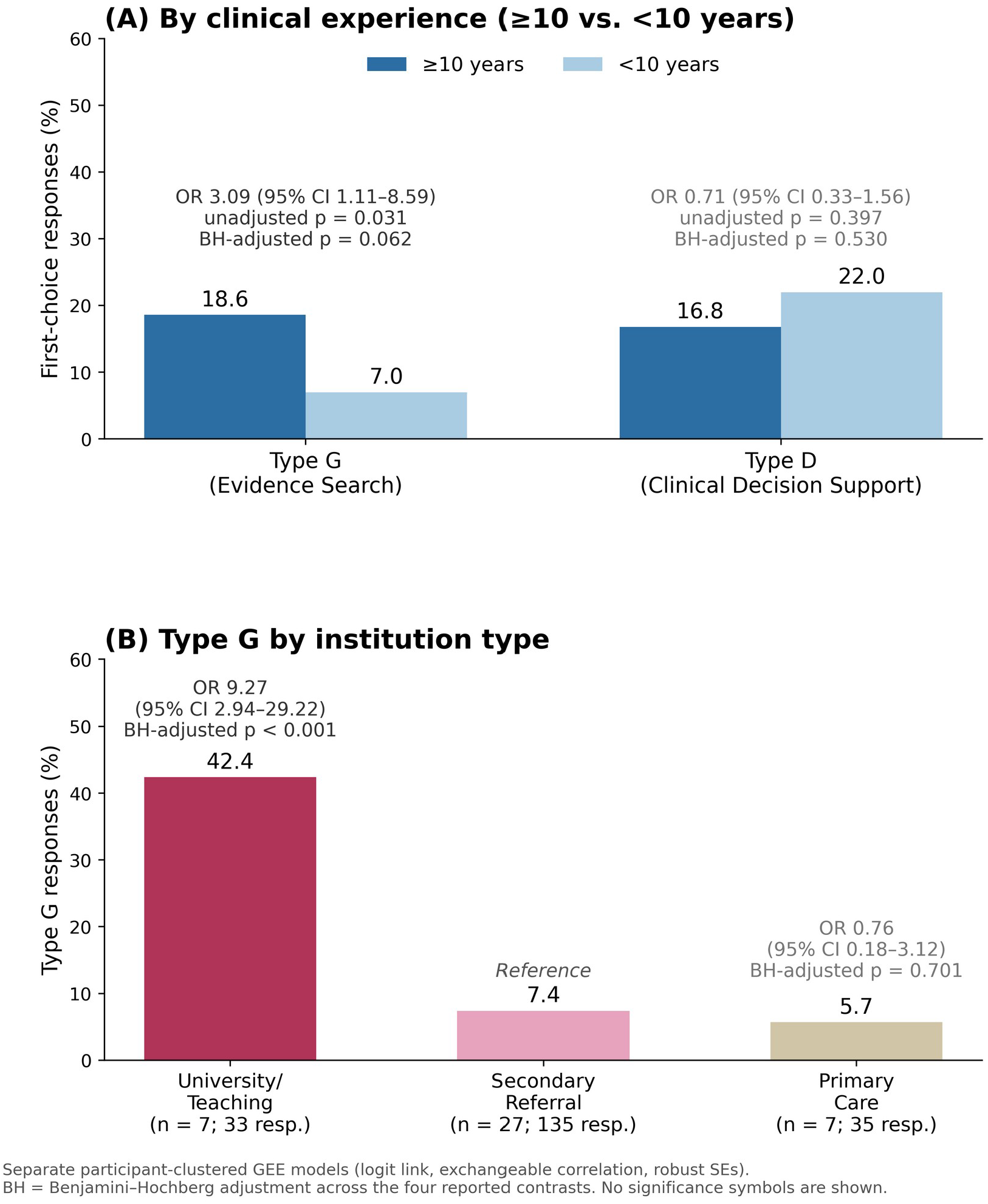
Subgroup Differences in AI Query Type Frequency by Clinical Experience and Institution Type. Values represent the percentage of first-choice responses within each group; effect estimates are from generalized estimating equations (GEE) with participant-level clustering. (A) Type G (Evidence Search) was selected more often by professionals with ≥10 years of experience (18.6% vs. 7.0%; OR 3.09, 95% CI 1.11–8.59; unadjusted p = 0.031; BH-adjusted p = 0.062) and Type D less often (16.8% vs. 22.0%; OR 0.71, 95% CI 0.33–1.56; BH-adjusted p = 0.530); neither experience contrast met the 0.05 threshold after adjustment. (B) Type G was selected more often in university/teaching hospitals (42.4%) than in secondary referral (7.4%, reference) and primary care (5.7%) settings (university/teaching vs. secondary referral: OR 9.27, 95% CI 2.94–29.22; BH-adjusted p < 0.001; primary care vs. secondary referral: OR 0.76, 95% CI 0.18–3.12; BH-adjusted p = 0.701). BH-adjusted p-values are corrected across the four reported contrasts; significance symbols are not used.

**Table 5.** First-Choice Query Type Frequency by Institution Type (Q1–Q5 Combined)

| **Query Type** | **Secondary Referral (n = 27, resp = 135)** | **University/Teaching (n = 7, resp = 33)** | **Primary Care (n = 7, resp = 35)** |
|---|---|---|---|
| **A Fact Check** | 18 (13.3%) | 1 (3.0%) | 7 (20.0%) |
| **B Differential Reasoning** | 43 (31.9%) | 9 (27.3%) | 9 (25.7%) |
| **C Treatment Protocol** | 25 (18.5%) | 4 (12.1%) | 6 (17.1%) |
| **D Clinical Decision Support** | 30 (22.2%) | 3 (9.1%) | 6 (17.1%) |
| **G Evidence Search** | 10 (7.4%) | 14 (42.4%) | 2 (5.7%) |
| **Other (E, F, H)** | 9 (6.7%) | 2 (6.1%) | 5 (14.3%) |

### Preferred Response Formats

Preferred response formats for the five representative high-frequency query types (A, B, C, D, G) varied considerably (Figure 5). For Type A (Fact Check), the format “value + normal range + interpretation” was preferred by 20 participants (46.5%). For Type B (Differential Reasoning), no single format predominated: preferences were split almost evenly across “prioritized differential list plus confirmatory tests” (15, 34.9%), “differential list with supporting evidence for each” (14, 32.6%), and “summary, evidence, values, synthesis, and next step” (13, 30.2%). For Type C (Treatment Protocol), “drug + dosage + precautions in table format” and “protocol + monitoring points” were tied as the most preferred formats (17/43, 39.5%, each); the remaining nine participants (20.9%) selected a step-by-step protocol. For Type D (Clinical Decision Support), a “pros-and-cons comparison table” was preferred by 27 participants (62.8%). For Type G (Evidence Search), the format “conclusion + source + evidence level (study design, sample size, reliability) + DOI” was preferred by 32 participants (74.4%), the highest single-format selection proportion among the five displayed query types.

**Figure 5.**
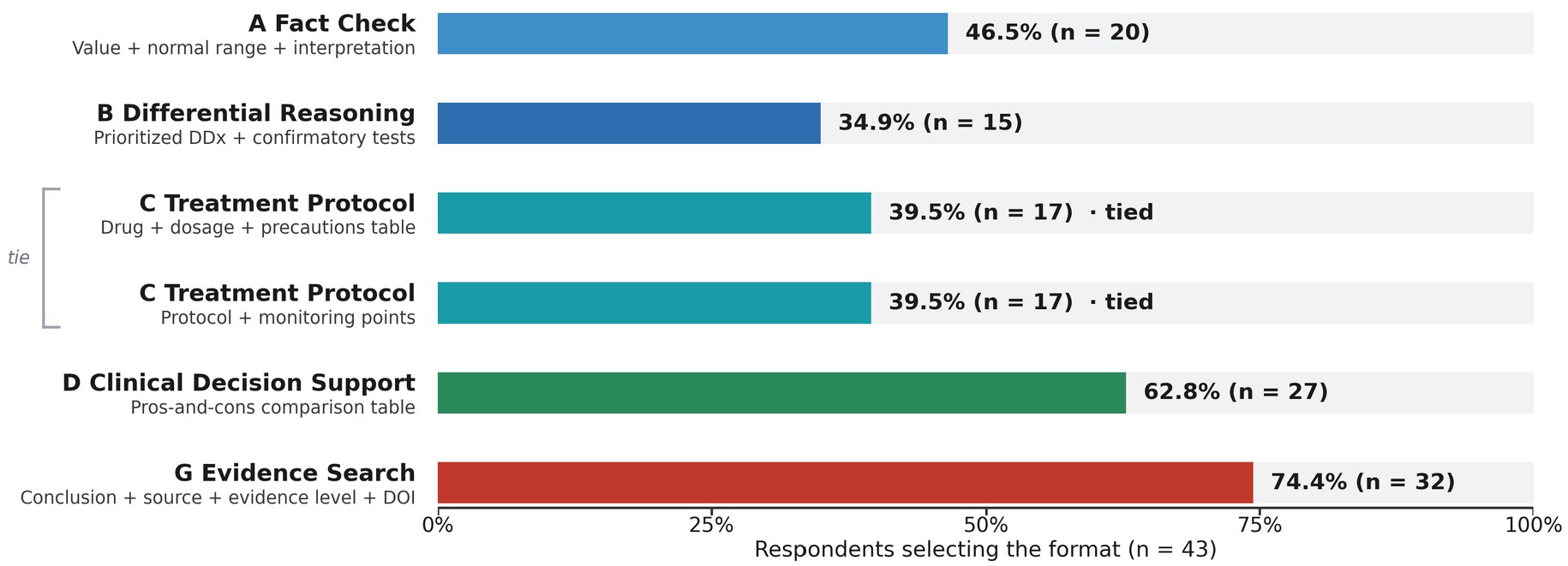
Preferred AI Response Formats by Query Type. Bars show the most frequently selected response formats for five representative query types (A, B, C, D, G), using all 43 respondents as the denominator. Both equally preferred Type C formats are shown (17/43, 39.5%, each). Type G (Evidence Search) had the largest single-format selection proportion among the displayed types (32/43, 74.4%). These proportions describe format preferences rather than formal consensus or taxonomy validity.

### Training Data Exclusion Criteria

In a multiple-selection item, respondents selected the following clinical-record types for exclusion from AI training: records of non-standard treatments performed at owner request (n = 29, 67.4%); records written by trainees or interns without attending verification (n = 26, 60.5%); records in which diagnosis was foregone due to owner financial constraints (n = 25, 58.1%); emergency records with incomplete or missing documentation (n = 24, 55.8%); and records using outdated treatment protocols inconsistent with current guidelines (n = 22, 51.2%).

### Essential Patient Context

In the multiple-selection item on essential patient information, species was selected by 41 of 43 respondents (95.3%), underlying diseases by 31 (72.1%), and current medications by 28 (65.1%). Age was selected by 15 (34.9%), breed by seven (16.3%), neuter status by four (9.3%), body weight by three (7.0%), owner financial circumstances/treatment willingness by one (2.3%), and other information by one (2.3%). These selections reflect respondents’ stated information priorities and do not establish the clinical necessity or sufficiency of individual variables.

## Discussion

### Interpretation of Core Findings

The taxonomy proposed here organizes veterinary AI queries into three categories and 21 subtypes, and the panel data show how first-choice preferences shifted across clinical stages. Type B (Differential Reasoning) was the most frequent first choice overall, suggesting that respondents prioritized clinical reasoning support over simple fact retrieval when asked about AI queries; whether this corresponds to actual query volume was not measured. This finding aligns with systematic evidence showing that clinical reasoning support and differential diagnosis are among the dominant use cases for LLMs in medical practice (Bedi et al., 2025; Qiu et al., 2025).

The frequent first-choice selection of Type D (Clinical Decision Support) for the post-consultation stage (50.0%) has not, to our knowledge, been reported previously. Respondents more often preferred to direct decision-support queries to AI after the consultation than during it; because this is a stated preference, it does not show when veterinarians actually use AI or why. If confirmed with usage data, this pattern would support evaluating structured post-consultation review functions alongside real-time support.

### Veterinary-Specific Characteristics

Several findings reflect veterinary-specific clinical characteristics. The high priority placed on species and underlying disease as essential patient context variables (95.3% and 72.1%, respectively) indicates that respondents prioritized species and comorbidity context. Current medications were also frequently selected (65.1%). In contrast, client communication support (Type F) was infrequently selected as the first choice (5/213 A–H responses), which contrasts descriptively with reports from human medicine in which patient communication is a common AI application (Ozkan et al., 2025). These rankings suggest that clinical reasoning and decision support were more often prioritized in this sample; low first-choice frequency does not establish low overall use or need for client communication support.

As noted above, existing veterinary AI research has focused on LLM performance on standardized examinations (Alonso Sousa et al., 2025) or general adoption surveys (Gabor et al., 2025), providing limited guidance on veterinary-specific query-type classification. The present taxonomy should be understood as a proposal developed from a preliminary review of real-world queries and literature-based refinement by the research team; the subsequent expert-panel survey characterized stage-specific preferences using that taxonomy. Formal confirmatory validation remains a necessary next step.

The present taxonomy shares structural similarities with, but is meaningfully differentiated from, existing classification frameworks in medical information science. Ely et al.’s seminal taxonomy of physician questions identified 69 generic question types centered on clinical management, drug information, and etiology (Ely et al., 1999); however, that framework predates LLM-based AI and does not address the AI-specific interaction modalities (e.g., prompt formulation, response format specification, hallucination risk) that characterize contemporary clinical AI use. More recently, Qiu et al. (Qiu et al., 2025) categorized physician LLM queries into broad functional domains (clinical support, documentation, research), but without the clinical-stage granularity or veterinary-specific subtypes provided by the present taxonomy. The distinguishing contribution of the current work is its dual-axis structure: query type classification is crossed with clinical stage, yielding a framework that describes which query types respondents reported preferring at each stage. It does not capture the reasons for, or the observed timing of, actual queries. Furthermore, the inclusion of a Terminology and Drug Reference category (Types M–U) reflects the species- and drug-specific lookup needs unique to multi-species veterinary practice, which are not separately distinguished in the human-medicine frameworks cited above.

### Differences by Experience and Institution Type

The separate GEE model suggested higher Type G selection among participants with ≥10 years of experience, but this association was sensitive to the multiple-testing family and additional adjustment. The four-contrast BH-adjusted p-value was 0.062, and the experience estimate was imprecise in same-sample and jointly adjusted analyses. Six of the seven university/teaching participants had ≥10 years of experience, underscoring overlap between experience and institution. The results therefore do not establish an independent experience effect. They may motivate larger studies of how clinical experience and institutional context relate to information needs (Ostropolets et al., 2020). The Type D experience contrast was not statistically significant.

The same analyses suggested that preference for Type G was elevated specifically in the university/teaching hospital group. No clear difference was detected between primary care and secondary referral settings, but that estimate was imprecise (OR 0.76, 95% CI 0.18–3.12) and does not show that the two settings are equivalent. This points to a concentration of evidence-search preference in the university/teaching group rather than a graded trend across institution types. One possible explanation, which this study did not measure, is the academic and research orientation of university/teaching hospitals; this interpretation is consistent with global surveys reporting that AI use for academic and research purposes exceeds clinical use in academic settings (Ozkan et al., 2025). The overlap between institution and experience noted above is another possible contributor. Because this estimate derives from a small subgroup (n = 7 participants), its magnitude should be regarded as preliminary.

### Implications for Veterinary AI Design

These findings are intended primarily for three audiences: developers of veterinary clinical AI systems, who may use the taxonomy to design stage-aware, context-sensitive response formats; researchers in veterinary informatics, who may use it as a scaffold for benchmark evaluation; and veterinary educators, who may use it to frame how AI tools are introduced across clinical training, including in resource-limited settings. For the educational audience, the query types are expressed in terms of what a clinician is trying to accomplish, which is also how competency frameworks describe professional practice. The nine domains of the AAVMC Competency-Based Veterinary Education framework (Chaney et al., 2024) offer a natural point of contact: Clinical Reasoning and Decision-making, Communication, and Scholarship each correspond to activities that recur among the query types reported here. Aligning the taxonomy with such a framework could allow AI-supported tasks to be situated within an existing curricular structure rather than treated as a separate skill set. We did not attempt that mapping, which would require validation of its own; we note it as a direction the present taxonomy makes available. The taxonomy is offered as a conceptual and evaluative framework rather than as a validation of any particular product; the authors’ affiliated system was used only as one source of preliminary query examples (Phase 1) and is neither evaluated nor promoted here. The 74.4% preference for “conclusion + source + evidence level + DOI” as the Type G response format suggests that respondents valued source attribution and evidence transparency. This finding supports considering evidence-level information and verifiable citations in veterinary AI response design. It does not establish that any particular response format improves clinical outcomes.

### Limitations

This study has several limitations that should be considered when interpreting its findings. First, the sample was drawn exclusively from South Korean veterinarians, and all chatbot log data originated from three animal hospitals in Ulsan and Gyeongsangnam-do (one secondary referral hospital, one cancer center, and one primary-care clinic). Veterinary practice norms, AI adoption rates, and clinical information needs vary substantially across countries and healthcare systems; accordingly, the taxonomy and reported preferences may not generalize to veterinarians in other regions, particularly those in North America, Europe, or lower-resource settings where AI tool availability differs markedly. Future multinational studies are needed to assess the cross-cultural robustness of the taxonomy. Second, participants were a self-selected convenience sample recruited through an open invitation in a single online community, so the panel is not a representative sample of the profession and the proportions reported here are not population estimates. The sample is skewed toward secondary referral hospitals (62.8%), and subgroup sizes for university/teaching hospitals (n = 7) and primary care clinics (n = 7) are insufficient for robust comparative inference; findings from these subgroups should be interpreted as preliminary and hypothesis-generating only. Third, the taxonomy was not formally validated. It was developed through a preliminary review of real-world queries and literature-based refinement by the research team, an approach consistent with exploratory taxonomy-generation methodology (Elo and Kyngäs, 2008), whereas the Phase 3 panel survey was used only to characterize query preferences and response requirements. The survey did not assess agreement with the subtype definitions or category assignments, and no inter-rater reliability metric (e.g., Cohen’s kappa) or content validity index (CVI) was obtained. Confirmatory studies applying formal reliability metrics to independent datasets are a necessary next step. The small number of participants in individual institution groups also limits the reliability of asymptotic GEE inference, and the joint sensitivity model has few outcome events relative to its number of coefficients. Fourth, stage-level rankings are self-reported preferences rather than observed query frequencies, and excluding two explicit non-use responses makes the A–H distributions conditional on selection of a listed query type; the survey did not measure reasons for querying, actual AI use, or clinical effects. Participants’ prior experience with AI or large language model tools was not recorded, so the preferences reported here may partly reflect differing familiarity with such systems. Fifth, the cross-sectional single-timepoint design cannot capture longitudinal changes in query preferences as AI capabilities and veterinary AI literacy continue to evolve rapidly. Sixth, the Type G and Type D outcomes were selected after the descriptive distributions of all eight types had been inspected, rather than pre-specified, and the planned permutation tests were not performed; the inferential results are therefore exploratory. Seventh, the coverage check reused the same queries that had informed the taxonomy and human review was limited to the non-converged cases, so it does not establish classification reliability. Future confirmatory studies with larger, nationally representative, multinational samples, formal reliability testing, and longitudinal designs are warranted to strengthen and extend these findings.

## Conclusions

This study proposes an exploratory taxonomy of veterinary AI query types comprising 3 categories and 21 subtypes. The taxonomy was developed from a preliminary review of real-world clinical query logs and refinement against the literature, and was then used in an expert panel study to elicit stage-specific query preferences from 43 veterinarians.

Three key findings emerge. First, first-choice query-type preferences differed descriptively by clinical stage, with Type B (Differential Reasoning) most frequently selected overall and Type D (Clinical Decision Support) most frequently selected for the post-consultation stage. Second, university/teaching affiliation was associated with Type G (Evidence Search) selection relative to secondary referral hospitals, while the experience contrast was sensitive to multiplicity correction and additional adjustment; small subgroup sizes limit inference. Third, for Type G (Evidence Search) 74.4% of respondents preferred a response format combining source attribution, evidence-level information, and a DOI link.

The taxonomy presented here provides a preliminary framework for the design of context-aware, stage-adaptive veterinary AI systems and benchmark evaluation frameworks. Future confirmatory validation using multinational, longitudinal datasets is warranted to strengthen and extend these findings.

## Author Contributions

Conceptualization, Hyeyeon Huh and Chan Huh; Methodology, Hyeyeon Huh; Software, Hyeyeon Huh; Formal analysis, Hyeyeon Huh; Investigation, Hyeyeon Huh and Chan Huh; Resources, Chan Huh, Jinsu Ahn and Minchul Park; Data curation, Hyeyeon Huh and Chan Huh; Writing — original draft preparation, Hyeyeon Huh; Writing — review and editing, Hyeyeon Huh, Chan Huh, Jinsu Ahn and Minchul Park; Visualization, Hyeyeon Huh; Project administration, Hyeyeon Huh and Chan Huh. All authors contributed to the article and approved the submitted version.

## Institutional Review Board Statement

The study protocol was approved through expedited review by the Institutional Review Board of Changwon National University, Republic of Korea, on 11 September 2026 (approval no. 7001066-202607-HR-053). The approved protocol described the prior use of 1,785 de-identified chatbot queries from 34 user accounts for questionnaire development and the subsequent anonymous expert-panel survey. All 43 survey responses analyzed in this study were collected after approval, with collection completed on 19 September 2026.

## Informed Consent Statement

Under the IRB-approved consent procedure, the full participant information sheet was presented on the first screen of the online survey. Participants were informed that voluntarily completing and submitting the questionnaire constituted consent to participate. No separate signed consent form or consent checkbox was used. Participants could discontinue the survey before submission without penalty. Because responses were anonymous, individual responses could not be identified for withdrawal after submission.

## Data Availability Statement

The survey data supporting the reported results are available from the corresponding author upon reasonable request. The classification prompts used for the post-taxonomy coverage check, with the run configuration (Supplementary Material S1), the taxonomy definitions supplied to the classifier (S2), the questionnaire as deployed with an English translation (S3), descriptive results for Modules 4–7 and the second-choice distributions (S4), the GEE analysis code with its output (S5), and a schematic of the deployed chatbot interface (S6) are provided as Supplementary Material; the full classification and de-identification pipeline code and the de-identified query dataset are available from the corresponding author on reasonable request.

## Acknowledgments

The authors thank the 43 veterinarians who participated in the survey, and the Choonok Company developer, who is not an author, who carried out the de-identification of the query logs. During the preparation of this manuscript, the authors used LLM-based tools — Claude (Anthropic; Claude Opus 4.1 and Claude Opus 5, accessed at claude.ai) and ChatGPT (OpenAI; the specific model version was not recorded, accessed at chatgpt.com) — to assist with manuscript preparation and with the analysis code. No text was accepted without author review, and no generative AI tool is listed as an author. The locally deployed model used to draft candidate query categories and to classify the query logs is described in the Methods. The authors have reviewed and edited all outputs and take full responsibility for the content of this publication.

