## Supplementary Material S1-S6 for "Development of an Exploratory Taxonomy of Veterinary AI Query Types and Stage-Specific Query Preferences: An Expert Panel Study"

*Source. The prompts below are the prompt strings of the classification pipeline (classify_pipeline.py and taxonomy.json), copied from the script that was run, and confirmed as such by the developer who ran it. They were written in Korean, which is the working language of the development team; the Korean text is given first, followed by an English translation prepared by the authors. Placeholders in braces are filled at run time. Section S1.6 additionally documents the serialized model-input format after application of the model-specific chat template.*

*Typesetting. Prompt lines longer than the page width are wrapped with a four-space continuation indent; no other change was made to the prompt text.*

*Scope. These prompts concern the coverage check in which the exploratory taxonomy was applied to the 1,785 de-identified user queries after the taxonomy had been established in Phase 2. They are not the prompts used for candidate generation in Phase 1, which were not retained.*

#### S1.1 Run configuration

| Item | Value |
|---|---|
| Model | nvidia/Gemma-4-26B-A4B-NVFP4 |
| Runtime | vLLM served locally (RTX 5090), called through LangChain (langchain-openai); the vLLM version was not recorded |
| Temperature | 0 |
| Input | All 1,785 de-identified user-authored queries (role = user); the output file contains 1,785 records with no duplicates |
| Run | 19 July 2026, 12:12-12:17 (KST) |
| Decision structure | Two stages: category, then subtype within the selected category |
| Decision readout | A single output token, read from the output-token log-probabilities |
| Verification | Binary Y/N suitability check against the assigned subtype definition |
| Reclassification | An unsuitable subtype was excluded and classification repeated, up to five iterations |
| Call granularity | One query per model call; no batching and no shared conversational context across queries |
| Non-clinical code | X (Other) |

#### S1.2 Category prompt

```
[system]
수의 임상 AI 챗봇에 수의사가 입력한 질문을 질문의 일차적 의도에 따라 분류한다.
1: Clinical Support - 눈앞의 임상 행위를 지원 (사실 확인, 감별진단, 치료 프로토콜, 입원/외래 판단,
    기록 작성, 보호자 설명, 근거 검색, 기전 학습)
2: Evidence-Based Research - 발표된 문헌·근거의 조회와 종합이 목적 (예후·생존기간 근거, 약물 근거,
    검사 해석 근거, 문헌 비교)
3: Terminology and Drug Reference - 특정 용어·질환·약물·시술 하나를 짚어 즉답을 구하는 조회 (용량,
    검사치 해석, 영상, 시술 단계, 질병 개요)
0: 기타 - 임상 질의가 아님 (잡담, 인사, 시스템 사용 문의)
숫자 하나만 출력한다.

[user]
{질문}
```

English translation:

```
[system]
Classify a question submitted by a veterinarian to a veterinary clinical AI chatbot according to
the primary intent of the question.
1: Clinical Support - supports an immediate clinical action (fact checking, differential diagnosis,
   treatment protocol, hospitalization/outpatient decision, record writing, owner explanation,
   evidence search, learning a mechanism)
2: Evidence-Based Research - the purpose is retrieval and synthesis of published literature and
   evidence (prognosis and survival-time evidence, drug evidence, evidence for test interpretation,
   literature comparison)
3: Terminology and Drug Reference - a lookup that points to one specific term, disease, drug, or
   procedure and seeks an immediate answer (dose, laboratory value interpretation, imaging,
   procedure steps, disease overview)
0: Other - not a clinical query (chitchat, greeting, question about using the system)
Output a single digit only.

[user]
{question}
```

Category decision read from the output-token log-probabilities:

| Output token | Category | Subtypes available at the second stage |
|---|---|---|
| 1 | Clinical Support Queries | A-H |
| 2 | Evidence-Based Research Queries | I-L |
| 3 | Terminology and Drug Reference Queries | M-U |
| 0 | Other | None; the query received the code X |

#### S1.3 Subtype prompts

One prompt per category. The definitions and examples listed in each prompt are those reproduced in Supplementary Material S2.

```
[system]
수의 임상 질문을 Clinical Support Queries 카테고리 내 세부 유형으로 분류한다.
A: Fact Check - Verification of factual clinical data; lab value accuracy, species-specific
    reference ranges (예: "What is the normal range for canine Lipase?")
B: Differential Reasoning - Multi-finding pattern synthesis for DDx generation (예: "BUN/Cre both
    elevated with K+ at 6.8 - what should I suspect?")
C: Treatment Protocol - Drug selection, dosing calculations, fluid therapy plans, or anesthetic
    protocols (예: "Fluid protocol for acute pancreatitis hospitalization?")
D: Clinical Decision - Hospitalization vs. outpatient judgment; triage and management decision
    support (예: "Hospitalization vs. outpatient follow-up for this CKD case?")
E: Documentation - SOAP note generation, discharge summaries, referral letter drafting (예: "Write a
    SOAP note for today's visit.")
F: Client Communication - Owner-facing explanations of diagnoses, treatments, or prognosis (예:
    "Draft a message explaining CKD diagnosis to the owner.")
G: Evidence Search - RCT lookup, clinical guideline retrieval, evidence-based practice queries (예:
    "Any RCTs comparing methimazole vs. radioactive iodine for feline hyperthyroidism?")
H: Education - Pathophysiology clarification, mechanism-of-action queries, self-directed learning
    (예: "Why are obese cats particularly susceptible to hepatic lipidosis?")
코드 문자 하나만 출력한다.

[user]
{질문}
[system]
수의 임상 질문을 Evidence-Based Research Queries 카테고리 내 세부 유형으로 분류한다.
I: Prognosis / Treatment Effect - Survival time, median survival, treatment outcome prediction (예:
    "MST for canine lymphoma?")
J: Drug / Protocol Evidence - Literature-based evidence for treatment choices, drug comparisons (예:
    "Cyclophosphamide use in feline mammary tumors?")
K: DDx / Test Interpretation - Literature-based test result context; evidence for differential
    diagnosis workup (예: "Isolated hypercholesterolemia differential?")
L: Literature Comparison - Comparative study retrieval; head-to-head comparison of interventions
    (예: "Clopidogrel vs. rivaroxaban for venous thrombosis?")
```

```
코드 문자 하나만 출력한다.

[user]
{질문}
[system]
수의 임상 질문을 Terminology and Drug Reference Queries 카테고리 내 세부 유형으로 분류한다.
M: Instant Answer - Single-term lookup initiated by a specific clinical term or acronym (예: "What
    is methylprednisolone?")
N: Clinical Approach - Standard clinical approach for a named condition or procedure (예:
    "Step-by-step approach for suspected acute hemorrhagic pancreatitis?")
O: DDx - Differential diagnosis list initiated by a single term with limited clinical context (예:
    "FNA differential diagnoses?")
P: Protocol - Standard treatment or diagnostic protocol for a named condition (예:
    "Methylprednisolone dosing protocol?")
Q: Lab Interpretation - Interpretation of a specific lab test result or panel (예: "What does
    elevated CBC mean?")
R: Dosage / Drug - Specific drug dosage, route, frequency, or drug-drug interaction lookup (예: "CRI
    dosage calculation?")
S: Imaging - Radiographic or ultrasound interpretation; imaging protocol lookup (예: "X-ray vs. CT -
    which should I order?")
T: Procedure - Step-by-step procedure guidance; surgical or diagnostic technique lookup (예: "Steps
    for thoracostomy tube placement?")
U: Disease / Diagnosis - Comprehensive disease overview initiated by a named condition (예:
    "Differential diagnoses for this tumor type?")
코드 문자 하나만 출력한다.

[user]
{질문}
```

English translation of the instruction lines, which are identical in the three prompts apart from the category name: “Classify the veterinary clinical question into a subtype within the <category> category.” and “Output a single code character only.” The subtype lines are already in English. 예 means “e.g.” and {질문} is the placeholder for the query.

#### S1.4 Retry message

When a verification step rejected an assignment, the next attempt appended the excluded codes to the user message, at both the category and the subtype stage:

```
[user]
{질문}

참고: 이전 시도에서 부적합 판정된 유형 코드 = [{제외 코드}]
```

English translation: “{question} / Note: subtype codes judged unsuitable in previous attempts = [{excluded codes}]”.

#### S1.5 Verification prompt

```
[system]
수의 임상 질문 분류를 검토한다. Y 또는 N 한 글자만 출력한다.

[user]
질문:
{질문}

부여된 유형: {코드} {유형명} - {정의}
이 유형이 질문에 명백히 맞지 않으면(예: 근거검색 질문에 단순조회 유형) N, 부합하거나 인접 유형과
    경합하는 정도면 Y.
```

English translation:

```
[system]
Review the classification of a veterinary clinical question. Output only one character, Y or N.

[user]
Question:
```

```
{question}

Assigned subtype: {code} {name} - {definition}
If this subtype clearly does not fit the question (for example, a simple-lookup subtype assigned
to an evidence-search question), answer N; if it fits, or merely competes with an adjacent
subtype, answer Y.
```

*Criterion actually applied. The retained verification prompt rejects an assignment only when the subtype clearly does not fit, and accepts it when the query merely competes with an adjacent subtype. This relaxed criterion was adopted after strict verification caused queries to cycle among the closely defined Evidence-Based Research subtypes (I-L). The prompt is not restricted to I-L: the relaxed criterion applies to every subtype. The earlier, stricter wording was not retained in the pipeline archive, so it cannot be reproduced here.*

*Operational consequence. A verification result of N removed the assigned subtype from the next classification attempt, and reclassification was repeated for up to five iterations. Queries that did not converge within five iterations (128 of 1,785; 7.2%) were independently adjudicated by all four authors. Only the final labels were retained, so the number of labels the authors changed was not recorded. The coverage-check labels were not used to estimate prevalence or frequency anywhere in the manuscript.*

#### S1.6 Assembled model input

The pipeline wrapped every prompt in the chat template of nvidia/Gemma-4-26B-A4B-NVFP4 before sending it. This section documents that serialized model-input format. Runtime query and exclusion-code values are represented here by the same placeholders used above; after substitution of those runtime values, this is the form sent to the model. The assembled input for the category prompt is shown first; the subtype and verification prompts were wrapped in the same way, with the bodies exactly as given in S1.3 and S1.5, so a reader can reconstruct them from this template. The thought channel was opened and closed empty, so generation resumed at the answer position and the decision could be read from the log-probabilities of a single output token.

```
<bos><|turn>system
[the system text of S1.2, unchanged]<turn|>
<|turn>user
{질문}<turn|>
<|turn>model
<|channel>thought
<channel|>
```

On a retry the user turn carried the exclusion note. Note that the excluded codes are rendered as a quoted list when the template is filled, so the assembled input reads:

```
<|turn>user
{질문}

참고: 이전 시도에서 부적합 판정된 유형 코드 = ['{제외 코드}']<turn|>
```

And the verification prompt, assembled in full:

```
<bos><|turn>system
수의 임상 질문 분류를 검토한다. Y 또는 N 한 글자만 출력한다.<turn|>
<|turn>user
질문:
{질문}

부여된 유형: {코드} {유형명} - {정의}
이 유형이 질문에 명백히 맞지 않으면(예: 근거검색 질문에 단순조회 유형) N, 부합하거나 인접 유형과
    경합하는 정도면 Y.<turn|>
<|turn>model
<|channel>thought
<channel|>
```

### Supplementary Material S2. Taxonomy Definitions and Examples Supplied to the Classifier

*Development of an Exploratory Taxonomy of Veterinary AI Query Types and Stage-Specific Query Preferences: An Expert Panel Study (Manuscript ID 1899253)*

*What this table reproduces. The definitions and illustrative examples below are taken from taxonomy.json and are those that were supplied to the classifier during the coverage check described in S1. The definitions are identical to those in Table 2 of the article. The examples are not: for seven subtypes (C, G, L, O, R, S, and U) the example in Table 2 was rewritten during revision so that readers can see where adjacent subtypes separate. Both versions are shown here, so that what the classifier received remains on the record. All examples are author-written illustrations, not verbatim logged queries. A separate non-clinical code, X (Other), was used for greetings, chitchat, system-use questions, and queries containing no clinical information request.*

| Code | Subtype name | Defining characteristic | Example supplied to the classifier | Example shown in Table 2 of the article |
|---|---|---|---|---|
| **Clinical Support Queries (A-H)** | | | | |
| A | Fact Check | Verification of factual clinical data; lab value accuracy, species-specific reference ranges | What is the normal range for canine Lipase? | (unchanged) |
| B | Differential Reasoning | Multi-finding pattern synthesis for DDx generation | BUN/Cre both elevated with K+ at 6.8 - what should I suspect? | (unchanged) |
| C | Treatment Protocol | Drug selection, dosing calculations, fluid therapy plans, or anesthetic protocols | Fluid protocol for acute pancreatitis hospitalization? | Fluid and analgesia plan for this 8-kg dog hospitalized with acute pancreatitis? |
| D | Clinical Decision | Hospitalization vs. outpatient judgment; triage and management decision support | Hospitalization vs. outpatient follow-up for this CKD case? | (unchanged) |
| E | Documentation | SOAP note generation, discharge summaries, referral letter drafting | Write a SOAP note for today's visit. | (unchanged) |
| F | Client Communication | Owner-facing explanations of diagnoses, treatments, or prognosis | Draft a message explaining CKD diagnosis to the owner. | (unchanged) |
| G | Evidence Search | RCT lookup, clinical guideline retrieval, evidence-based practice queries | Any RCTs comparing methimazole vs. radioactive iodine for feline hyperthyroidism? | Is there a current guideline for managing hyperthyroidism in a cat that also has CKD? I need it for this patient's plan. |
| H | Education | Pathophysiology clarification, mechanism-of-action queries, self-directed learning | Why are obese cats particularly susceptible to hepatic lipidosis? | (unchanged) |
| **Evidence-Based Research Queries (I-L)** | | | | |
| I | Prognosis / Treatment Effect | Survival time, median survival, treatment outcome prediction | MST for canine lymphoma? | (unchanged) |

| Code | Subtype name | Defining characteristic | Example supplied to the classifier | Example shown in Table 2 of the article |
|---|---|---|---|---|
| J | Drug / Protocol Evidence | Literature-based evidence for treatment choices, drug comparisons | Cyclophosphamide use in feline mammary tumors? | (unchanged) |
| K | DDx / Test Interpretation | Literature-based test result context; evidence for differential diagnosis workup | Isolated hypercholesterolemia differential? | (unchanged) |
| L | Literature Comparison | Comparative study retrieval; head-to-head comparison of interventions | Clopidogrel vs. rivaroxaban for venous thrombosis? | What do published studies comparing clopidogrel and rivaroxaban for thromboprophylaxis in dogs report? |
| **Terminology and Drug Reference Queries (M-U)** | | | | |
| M | Instant Answer | Single-term lookup initiated by a specific clinical term or acronym | What is methylprednisolone? | (unchanged) |
| N | Clinical Approach | Standard clinical approach for a named condition or procedure | Step-by-step approach for suspected acute hemorrhagic pancreatitis? | (unchanged) |
| O | DDx | Differential diagnosis list initiated by a single term with limited clinical context | FNA differential diagnoses? | Differentials for a round-cell population on a skin FNA? |
| P | Protocol | Standard treatment or diagnostic protocol for a named condition | Methylprednisolone dosing protocol? | (unchanged) |
| Q | Lab Interpretation | Interpretation of a specific lab test result or panel | What does elevated CBC mean? | (unchanged) |
| R | Dosage / Drug | Specific drug dosage, route, frequency, or drug-drug interaction lookup | CRI dosage calculation? | Fentanyl CRI dose range for dogs? |
| S | Imaging | Radiographic or ultrasound interpretation; imaging protocol lookup | X-ray vs. CT - which should I order? | What does a diffuse unstructured interstitial pattern on thoracic radiographs indicate? |
| T | Procedure | Step-by-step procedure guidance; surgical or diagnostic technique lookup | Steps for thoracostomy tube placement? | (unchanged) |
| U | Disease / Diagnosis | Comprehensive disease overview initiated by a named condition | Differential diagnoses for this tumor type? | Overview of canine hemangiosarcoma: diagnosis, staging, and prognosis? |

Primary-intent rule. Each query received one subtype according to its primary information-seeking intent. Types A-H represent point-of-care clinical support, Types I-L represent evidence retrieval and appraisal, and Types M-U represent terminology, drug, test, imaging, procedure, or disease reference entry points. No separate coding manual, list of borderline cases, or tie-break hierarchy was used during classification.

### Supplementary Material S3. Survey Questionnaire (as deployed)

*Manuscript 1899253 — Development of an Exploratory Taxonomy of Veterinary AI Query Types and Stage-Specific Query Preferences: An Expert Panel Study*

##### Notes on this version

Source. The questionnaire below reproduces the Google Forms version reviewed and approved by the Institutional Review Board of Changwon National University (approval no. 7001066-202607-HR-053, 11 September 2026) and deployed for data collection. The reproduced items were checked against the form and the exported response data (collection completed on 19 September 2026). Part A is an English translation prepared by the authors; Part B is the Korean original. Links to the online forms have been removed.

Response options. In this Supplementary Material, the response options are reproduced as they appeared in the deployed online form, based on the form and the exported response data. All eight Clinical Support query types (A–H) were available as response options for each clinical stage; the legend listing the types was shown above the stage questions. An “Other (please specify)” option was offered at Q1 and Q4, and two respondents used it.

Coding of background responses. Clinical experience was reported as free text and was converted to years: entries giving months were divided by 12, bare numeric entries were read as years, and two entries written in Korean (“10 냔” and “7 녀차”) were read as 10 years and 7 years respectively. For clinical specialty, the internal-medicine grouping reported in the article (n = 26) combines general and preventive medicine, dermatology-associated, gastrointestinal and endocrine, and oncology-related internal-medicine responses; emergency and critical care was offered as a single combined option in the questionnaire.

Items analyzed in the article. The main text reports the first-choice responses to Module 1 Q1–Q5, the response-format items Module 1 Q7–Q10 and Q13, Module 2 Q1, and Module 3 Q1. Descriptive results for the other fixed-choice items are given in Supplementary Material S4. Free-text items were not coded or analyzed.

#### Part A. English translation

##### Participant information (first screen of the survey)

Study title: An exploratory typology of veterinarians' AI query patterns by clinical stage: an expert panel study

This is an exploratory study that aims to classify the types of questions veterinarians ask artificial intelligence (AI) chatbots in clinical practice according to clinical stage, experience, and institution type. Before deciding whether to take part, please read the following explanation carefully. The study is conducted only with those who voluntarily indicate their willingness to participate.

※ Please do not enter, in free-text items, any information that could identify a specific person or case, such as the name or contact details of a patient's owner, a hospital name, or a chart number.

1. Background and purpose. The clinical use of AI chatbots in veterinary medicine is increasing, but there is little systematic classification of the types of questions veterinarians ask at each clinical stage. This study aims to develop an exploratory framework for classifying veterinarians' AI queries through an expert survey.

2. Who can take part. About 40 veterinarians (minimum 30, maximum 60) who hold a veterinary license and are engaged in clinical practice at the time of the survey. Those without a license or not engaged in clinical practice are not eligible.

3. What participation involves. If you agree, you will complete an anonymous online survey on the AI query types you prefer (first and second choice) at five clinical stages (before consultation, before tests, after tests, immediately after consultation, and outside working hours), preferred answer

formats for each type, patient information needed in AI answers, types of records that should be excluded from AI training, and the situations in which you would use research/evidence queries and terminology/drug reference queries.

4. Duration. A single response taking about 20–30 minutes (survey period: within one month of IRB approval).

5. Withdrawal. You may stop at any time before submitting the survey without any disadvantage. Incomplete responses are excluded from analysis. After submission, however, responses are stored as anonymous data that are not linked to identifying information, so it is not possible to identify and withdraw or delete your response.

6. Risks. This is a non-invasive, anonymous survey. No risk beyond that ordinarily encountered in daily life is expected, and the risk of participation is minimal.

7. Benefits. There is no direct medical benefit to you. As a token of thanks, those who complete the survey will receive a mobile gift voucher worth KRW 30,000. If you wish to receive it, please leave your mobile phone number through the separate link on the last screen of the survey; this information is not linked to your survey responses and is destroyed immediately after the voucher is sent. Not applying for the voucher carries no disadvantage, and your responses are analyzed in the same way. Your responses will contribute to a better understanding of how veterinary AI systems should be designed.

8. Not participating. You are free not to take part, and there is no disadvantage if you do not.

9. Privacy and confidentiality. This anonymous survey does not collect personally identifying information such as name, contact details, or IP address, nor sensitive information under Article 23 of the Personal Information Protection Act. Contact details for the voucher are collected through a separate route, are not linked to survey responses in any way, and are destroyed immediately after sending. Responses are processed only statistically, and no identifying information is used in journal or conference publications. The Institutional Review Board of Changwon National University may inspect study data within the scope of applicable regulations. Study data are retained for three years after the end of the study and then destroyed.

10. Disclosure of interests. The preliminary items of this study were derived by reviewing de-identified query logs from a pilot chatbot operated by Choonok Co., Ltd., and the investigators are affiliated with that company. There is no separate research funding for this study; only the cost of participant vouchers is paid from the TIPS program (Ministry of SMEs and Startups) carried out by the company. The related conflict of interest has been reported to the Institutional Review Board of Changwon National University.

11. Contact. Principal investigator: Hyeyeon Huh /

※ Please respond only once. ※ This study has been reviewed by the Institutional Review Board of Changwon National University (approval no. 7001066-202607-HR-053). ※ Responding to and submitting this survey is considered consent to participate in the study. If you do not wish to participate, please close this window. Thank you for your time.

##### Eligibility

**S1. Do you hold a veterinary license?**

☐ Yes
☐ No

**S2. Are you currently engaged in clinical practice?**

☐ Yes
☐ No

*If "No" is selected for either question, the respondent is not eligible and the survey ends. Employees of Choonok Co., Ltd. may not participate.*

#### Background information

**1. Type of institution**

- ☐ Primary care clinic
- ☐ Secondary referral hospital
- ☐ University/teaching hospital
- ☐ Research institution
- ☐ Other (________)

**2. Clinical experience: about ____ years ____ months** *[free text]*

**3. Main clinical specialty (multiple entries allowed): ________** *[free text]*

*Query-type legend shown to respondents. Clinical (A–H): A fact check · B differential/reasoning · C treatment/protocol · D clinical decision support · E chart/document writing · F client communication · G literature/evidence search · H education/explanation. Research (I–L): I prognosis/treatment effect · J drug/procedure evidence · K differential/test interpretation · L literature comparison. Terminology/drug (M–U): M instant answer · N clinical approach · O differential diagnosis · P protocol · Q test interpretation · R dose/administration · S imaging interpretation · T procedure · U disease/diagnosis.*

#### Module 1. Clinical queries — query types by stage and preferred answer structure

**Q1. Before consultation — Which type of question would you ask AI? (1st choice ____ 2nd choice ____; enter option numbers)** *[ranked]*

- ☐ 1) Quick fact check (Type A)
- ☐ 2) Differential reasoning (Type B)
- ☐ 3) Treatment/protocol (Type C)
- ☐ 4) Clinical decision support (Type D)
- ☐ 5) Chart/document writing (Type E)
- ☐ 6) Client communication (Type F)
- ☐ 7) Literature/evidence search (Type G)
- ☐ 8) Education/explanation request (Type H)
- ☐ 9) Other (please specify)

**Q2. During consultation (before tests) — Which type of question would you ask AI? (1st choice ____ 2nd choice ____; enter option numbers)** *[ranked]*

- ☐ 1) Quick fact check (Type A)
- ☐ 2) Differential reasoning (Type B)
- ☐ 3) Treatment/protocol (Type C)
- ☐ 4) Clinical decision support (Type D)
- ☐ 5) Chart/document writing (Type E)
- ☐ 6) Client communication (Type F)
- ☐ 7) Literature/evidence search (Type G)
- ☐ 8) Education/explanation request (Type H)

**Q3. During consultation (after tests) — Which type of question would you ask AI? (1st choice ____ 2nd choice ____; enter option numbers)** *[ranked]*

- ☐ 1) Quick fact check (Type A)
- ☐ 2) Differential reasoning (Type B)
- ☐ 3) Treatment/protocol (Type C)
- ☐ 4) Clinical decision support (Type D)
- ☐ 5) Chart/document writing (Type E)
- ☐ 6) Client communication (Type F)
- ☐ 7) Literature/evidence search (Type G)
- ☐ 8) Education/explanation request (Type H)

**Q4. Immediately after consultation — Which type of question would you ask AI? (1st choice ____ 2nd choice ____; enter option numbers)** *[ranked]*

- ☐ 1) Quick fact check (Type A)
- ☐ 2) Differential reasoning (Type B)
- ☐ 3) Treatment/protocol (Type C)
- ☐ 4) Clinical decision support (Type D)
- ☐ 5) Chart/document writing (Type E)
- ☐ 6) Client communication (Type F)
- ☐ 7) Literature/evidence search (Type G)
- ☐ 8) Education/explanation request (Type H)
- ☐ 9) Other (please specify)

**Q5. Outside working hours — Which type of question would you ask AI? (1st choice ____ 2nd choice ____; enter option numbers)** *[ranked]*

- ☐ 1) Quick fact check (Type A)
- ☐ 2) Differential reasoning (Type B)
- ☐ 3) Treatment/protocol (Type C)
- ☐ 4) Clinical decision support (Type D)
- ☐ 5) Chart/document writing (Type E)
- ☐ 6) Client communication (Type F)
- ☐ 7) Literature/evidence search (Type G)
- ☐ 8) Education/explanation request (Type H)

**Q6. Examples of actual clinical questions** *[free text]*

**Q7. Answer structure for Type A (fact check) (choose one)**

- ☐ ① Key values only, concisely
- ☐ ② Values + brief clinical context
- ☐ ③ Values + comparison with normal range + interpretation
- ☐ ④ Other

**Q8. Answer structure for Type B (differential/reasoning) (choose one)**

- ☐ ① DDx list + supporting evidence for each
- ☐ ② Summary → evidence → values → synthesis → next step
- ☐ ③ Prioritized DDx + confirmatory tests
- ☐ ④ Other

**Q9. Answer structure for Type C (treatment/protocol) (choose one)**

- ☐ ① Step-by-step protocol
- ☐ ② Table of drug name + dose + precautions
- ☐ ③ Protocol + monitoring points

**Q10. Answer structure for Type D (decision support) (choose one)**

- ☐ ① Pros-and-cons comparison table
- ☐ ② Conclusion first, evidence after

**Q11. Answer structure for Type E (chart writing) (choose one)**

- ☐ ① Basic S/O/A/P structure only
- ☐ ② S/O/A/P + English–Korean terms + three-part Plan
- ☐ ③ S/O/A/P + annotations explaining the reasoning
- ☐ ④ Other

**Q12. Answer structure for Type F (client guidance) (choose one)**

- ☐ ① Short message — 2–3 lines, key points only (messenger/text)
- ☐ ② Medium explanation — 5–6 lines, key point + reason + precautions
- ☐ ③ Detailed guidance — including precautions, schedule, and how to give medication

**Q13. Answer structure for Type G (evidence search) (choose one)**

☐ ① Conclusion + DOI link only
☐ ② Conclusion + source (title, authors, year) + DOI link
☐ ③ Conclusion + source + level of evidence (study design, number of cases, reliability) + DOI link

**Q14. Answer structure for Type H (education/explanation) (choose one)**

☐ ① Core mechanism in 3–5 lines
☐ ② Mechanism + clinical link
☐ ③ Textbook-level detail

**Q15. Elements that must be included in AI answers** *[free text]*

##### Module 2. Clinical queries — essential patient information

**Q1. Patient information that must be reflected in AI answers (select all that apply)** *[multiple selection]*

☐ ① Species (dog/cat)
☐ ② Breed
☐ ③ Age
☐ ④ Body weight
☐ ⑤ Underlying disease (existing diagnoses)
☐ ⑥ Current medications
☐ ⑦ Neuter status
☐ ⑧ Owner's financial situation/willingness to treat
☐ ⑨ Other (please specify)

**Q2. Patient information that would be good to have (rank in order of priority: 1, 2, 3 …)** *[ranked]*

☐ ① Species (dog/cat)
☐ ② Breed
☐ ③ Age
☐ ④ Body weight
☐ ⑤ Underlying disease (existing diagnoses)
☐ ⑥ Current medications
☐ ⑦ Neuter status
☐ ⑧ Owner's financial situation/willingness to treat
☐ ⑨ Other (please specify)

**Q3. Experiences in which contextual information changed your judgment** *[free text]*

##### Module 3. Clinical queries — records to exclude from training and bias prevention

**Q1. Types of clinical records that should be excluded from AI training (select all that apply)** *[multiple selection]*

☐ ① Records in which the owner declined tests for financial reasons, so only symptomatic treatment was given without a diagnosis
☐ ② Emergency records that are incomplete or have many missing items
☐ ③ Records of treatment with outdated protocols (differing from current guidelines)
☐ ④ Records written by trainees/interns without attending verification
☐ ⑤ Records of non-standard treatment given at the owner's request
☐ ⑥ Other (please specify)

**Q2. Methods for preventing data bias (select all that apply)** *[multiple selection]*

☐ ① Secure data evenly from hospitals in various regions (Seoul/provinces)
☐ ② Collect data from primary and secondary/tertiary hospitals in balance
☐ ③ Deliberately supplement data on rare breeds, not only popular breeds
☐ ④ Secure data from specialties such as surgery, ophthalmology, and dermatology, not only internal medicine
☐ ⑤ Supplement gaps in domestic data with international papers/guidelines
☐ ⑥ Other (please specify)

##### Module 4. Research/evidence queries — situations of use and answer priorities

**Q1. Situations in which you would use Type I (prognosis/treatment effect) (1st ____ 2nd ____)** *[ranked]*

☐ ① When an owner asks, “How long will my pet live?”
☐ ② When comparing success rates/survival times while planning treatment
☐ ③ When reassessing prognosis because the patient has deteriorated
☐ ④ When checking recurrence rates or long-term prognosis after surgery

**Q2. Situations in which you would use Type J (drug/procedure) (1st ____ 2nd ____)** *[ranked]*

☐ ① When checking the dose and administration of a drug used for the first time
☐ ② When checking whether the recommended dose of a familiar drug has changed
☐ ③ When looking for an alternative drug because of adverse effects
☐ ④ When wondering about interactions between two drugs used together
☐ ⑤ Other (please specify)

**Q3. Situations in which you would use Type K (differential/test) (1st ____ 2nd ____)** *[ranked]*

☐ ① When test values are equivocal and need interpretation
☐ ② When atypical signs require broadening the DDx
☐ ③ When imaging findings point to several diagnoses
☐ ④ When a rare disease is suspected and differential criteria need checking
☐ ⑤ Other (please specify)

**Q4. Situations in which you would use Type L (literature comparison) (1st ____ 2nd ____)** *[ranked]*

☐ ① When checking the evidence on whether treatment A or B is better
☐ ② When checking the evidence for a treatment the owner saw on the internet
☐ ③ When looking for evidence on an approach heard at a conference/seminar
☐ ④ When evidence is needed to decide whether to change an existing protocol
☐ ⑤ Other (please specify)

**Q5. Additional question types beyond Types I–L** *[free text]*

**Q6. Priority of information in answers citing papers (rank 1, 2, 3 …)** *[ranked]*

☐ ① Basic paper information (authors, journal, year)
☐ ② Conclusion/key figures (survival time, success rate, etc.)
☐ ③ Study population (species, breed, number of animals)
☐ ④ Clinical applicability (usable in the domestic setting)

**Q7. How should the level of evidence be expressed? (choose one)**

☐ A — State each source (title, authors, year)
☐ B — State the size of the study (e.g., 1 RCT, cohort of 138 animals)
☐ C — Show reliability as a grade (strong/moderate/weak evidence)
☐ D — Other (please write)

**Q8. Where should the level of evidence appear? (choose one)**

☐ A — Before the answer
☐ B — After the answer
☐ C — Naturally within the sentences
☐ D — Other (please write)

**Q9. Structure of answers based on research evidence (choose one)**

☐ A — Conclusion first
☐ B — PICO order
☐ C — Comparison table
☐ D — Other (please write)

**Q10. Things that must not be done when citing papers** *[free text]*

**Q11. Elements that must be included in answers citing papers (select all that apply)** *[multiple selection]*

☐ ① Source (title, authors, year)
☐ ② Number of cases (n)
☐ ③ Species/breed studied
☐ ④ Limitations of the study
☐ ⑤ Applicability in Korea

#### Module 5. Research/evidence queries — learning points and answer order

**Q1. Clinical research — learning points (rank 1, 2, 3 …)** *[ranked]*

☐ ① Summary of aim + methods
☐ ② Study population (species, breed, n)
☐ ③ Result values (P-value, survival time, etc.)
☐ ④ Conclusion

**Q2. Drug/procedure research — learning points (rank)** *[ranked]*

☐ ① Drug name (INN) + dose + administration
☐ ② Main adverse effects + monitoring items
☐ ③ Study population (species, breed, n)
☐ ④ Conclusion + clinical recommendations

**Q3. Case reports — learning points (rank)** *[ranked]*

☐ ① Patient signalment (species, breed, age)
☐ ② Main clinical findings + course
☐ ③ Treatment + outcome
☐ ④ Number of cases + limits to generalization

**Q4. Basic/conceptual research — learning points (rank)** *[ranked]*

☐ ① Explanation of the key mechanism/pathway
☐ ② Clinical applicability
☐ ③ Conclusion + future research directions
☐ ④ List of key prior studies cited

**Q5. Answer order for Type I (rank)** *[ranked]*

☐ ① Survival time/success rate
☐ ② P-value or statistical significance
☐ ③ Difference between groups (treatment vs. control)
☐ ④ Number of cases (n) + study design
☐ ⑤ Limits to clinical application (e.g., single case)
☐ ⑥ Other (please specify)

**Q6. Answer order for Type J (rank)** *[ranked]*

☐ ① Drug name (INN) + specific dose
☐ ② Route + dosing interval
☐ ③ List of main adverse effects
☐ ④ Monitoring items (what to check and when)
☐ ⑤ Contraindications/precautions

**Q7. Answer order for Type K (rank)** *[ranked]*

☐ ① Differential diagnosis list (in order of likelihood)
☐ ② Distinguishing features of each diagnosis
☐ ③ How to confirm (recommended tests)
☐ ④ Keep English terms (ddx, head tilt, etc.)
☐ ⑤ Indicate urgency/diagnoses to rule out first

**Q8. Answer order for Type L (rank)** *[ranked]*

☐ ① Pros and cons of each option
☐ ② Comparison table (A vs. B side by side)
☐ ③ Source studies for the evidence
☐ ④ Comparison of the level of evidence (how reliable the studies are)
☐ ⑤ Recommendations by clinical situation
☐ ⑥ Other (please specify)

**Q9. Length/form of summaries (choose one)**

☐ A — Key point only, in 1–2 sentences
☐ B — Conclusion + figures + limitations, in 3–5 sentences
☐ C — Divided into sections
☐ D — Other (please write)

**Q10. Elements that must be included in summaries** *[free text]*

**Q11. Additional comments on research/evidence queries** *[free text]*

##### Module 6. Research/evidence queries — research to exclude from training

**Q1. Types of research that should be excluded from AI training** *[free text]*

**Q2. If you have had difficulty applying international guidelines in Korea, please describe your experience.** *[free text]*

##### Module 7. Terminology/drug reference queries — situations of use and answer order

**Q1. Situations in which you would use Type M (instant answer) (1st ____ 2nd ____)** *[ranked]*

☐ ① When an unfamiliar term comes up during a consultation
☐ ② When checking quickly while reading a paper or textbook
☐ ③ When explaining to a junior colleague/student
☐ ④ Other (please specify)

**Q2. Type N (clinical approach) (1st ____ 2nd ____)** *[ranked]*

☐ ① When a patient with an unfamiliar disease/condition presents
☐ ② When checking the work-up order because it is unclear
☐ ③ When quickly checking the initial approach in an emergency
☐ ④ When an approach from another field (ophthalmology, orthopedics, etc.) is needed

**Q3. Type O (differential diagnosis) (1st ____ 2nd ____)** *[ranked]*

☐ ① When checking that no differential has been missed
☐ ② When test results point to several diagnoses
☐ ③ When explaining the list of possible diagnoses to an owner
☐ ④ When wanting to consider uncommon differentials broadly

**Q4. Type P (protocol) (1st ____ 2nd ____)** *[ranked]*

☐ ① When checking the exact dose/interval just before prescribing
☐ ② When organizing the order or timing of combined treatments
☐ ③ When dose adjustment is needed in special patients (renal failure, liver disease, etc.)
☐ ④ When setting up a monitoring protocol for inpatients
☐ ⑤ Other (please specify)

**Q5. Type Q (test interpretation) (1st ____ 2nd ____)** *[ranked]*

☐ ① When checking an unfamiliar item on a test report
☐ ② When values are equivocal and normal/abnormal is hard to judge
☐ ③ When several values must be interpreted together
☐ ④ Other (please specify)

**Q6. Type R (dose/administration) (1st ____ 2nd ____)** *[ranked]*

☐ ① When checking the dose of a drug used for the first time
☐ ② When the dosing interval (q8h, q12h, etc.) is unclear
☐ ③ When calculations for special administration such as CRI are needed
☐ ④ Other (please specify)

**Q7. Type S (imaging interpretation) (1st ____ 2nd ____)** *[ranked]*

☐ ① When imaging findings need to be interpreted
☐ ② When deciding which imaging test to choose (X-ray vs. ultrasound vs. CT, etc.)
☐ ③ When normal vs. abnormal must be distinguished on imaging
☐ ④ Other (please specify)

**Q8. Type T (procedure) (1st ____ 2nd ____)** *[ranked]*

☐ ① When the order or method of a procedure is unclear
☐ ② When checking the steps of a procedure performed for the first time
☐ ③ When checking how to manage complications during a procedure
☐ ④ Other (please specify)

**Q9. Type U (disease/diagnosis) (1st ____ 2nd ____)** *[ranked]*

☐ ① When a rare disease is suspected
☐ ② When wanting to check diagnostic criteria exactly
☐ ③ When prognosis must be explained to an owner
☐ ④ Other (please specify)

**Q10. Answer order for Type M (rank)** *[ranked]*

☐ ① Definition (what it is)
☐ ② Normal range/reference value
☐ ③ Clinical significance (why it matters)
☐ ④ Precautions

**Q11. Answer order for Type N (rank)** *[ranked]*

☐ ① Initial approach order (work-up)
☐ ② Essential tests
☐ ③ Differential diagnosis list
☐ ④ Criteria for judging urgency

**Q12. Answer order for Type O (rank)** *[ranked]*

☐ ① Differential diagnosis list (in order of likelihood)
☐ ② Distinguishing points of each diagnosis
☐ ③ Confirmatory tests
☐ ④ Diagnoses to rule out urgently

**Q13. Answer order for Type P (rank)** *[ranked]*

☐ ① Drug name + dose
☐ ② Route and interval
☐ ③ Monitoring points
☐ ④ Contraindications/precautions

**Q14. Answer order for Type Q (rank)** *[ranked]*

☐ ① Normal range
☐ ② Clinical significance of abnormality
☐ ③ Related differential diagnoses
☐ ④ Recommended additional tests

**Q15. Answer order for Type R (rank)** *[ranked]*

☐ ① Dose (mg/kg)
☐ ② Route and interval
☐ ③ Criteria for dose adjustment (renal failure, etc.)
☐ ④ Adverse effects/monitoring

**Q16. Answer order for Type S (rank)** *[ranked]*

☐ ① Description of findings
☐ ② Differential diagnoses
☐ ③ Normal vs. abnormal
☐ ④ Additional tests/follow-up recommendations

**Q17. Answer order for Type T (rank)** *[ranked]*

☐ ① Procedure order/method
☐ ② Supplies/equipment
☐ ③ Management of complications
☐ ④ Precautions/contraindications

**Q18. Answer order for Type U (rank)** *[ranked]*

☐ ① Definition/diagnostic criteria
☐ ② Differential diagnoses
☐ ③ Prognosis/course
☐ ④ Overview of treatment options
☐ ⑤ Other (please specify)

**Q19. Overall answer style for terminology queries (choose one)**

☐ A — Prose (written in sentences)
☐ B — Structured ([Definition][Dose][Precautions] labels)
☐ C — Other (please write)

**Q20. Additional comments** *[free text]*

##### Closing screen

The survey is complete. Thank you for participating. As a token of thanks, we will send a mobile gift voucher worth KRW 30,000. If you wish to receive it, please leave your mobile phone number via the link below (a separate form, not linked to this survey). The number you enter is not linked to your survey responses and is destroyed immediately after the voucher is sent. Your responses are used in the study in the same way whether or not you apply, without any disadvantage.

#### Part B. Korean original (국문 원문)

### 연구 참여 설명 (설문 첫 화면)

연구과제명: 임상 단계별 수의사의 AI 질의 패턴에 대한 탐색적 유형화: 전문가 패널 연구

본 연구는 수의 임상현장에서 수의사가 인공지능(AI) 챗봇에 하는 질의의 유형을 임상 단계·경력·기관유형에 따라 분류하기 위한 탐색적 연구입니다. 귀하는 본 연구에 참여할 것인지 결정하기 전에 아래 설명을 신중히 읽어보시기 바랍니다. 본 연구는 자발적으로 참여 의사를 밝히신 분에 한하여 수행됩니다.

※ 서술형 문항에는 환자·보호자의 성명, 연락처, 병원명, 차트번호 등 특정 개인이나 진료 사례를 식별할 수 있는 정보를 기재하지 말아 주십시오.

1. 연구의 배경과 목적: AI 챗봇의 수의 임상 활용이 늘고 있으나, 수의사가 임상 단계별로 어떤 유형의 질의를 하는지에 대한 체계적 분류는 부족합니다. 본 연구는 전문가 설문을 통해 수의사의 AI 질의 유형을 분류하는 탐색적 체계를 개발하고자 합니다.

2. 연구 참여 대상: 수의사 면허를 보유하고 조사 시점에 임상 진료에 종사하는 수의사 약 40 명(최소 30 명 ~ 최대 60 명)이 참여합니다. 수의사 면허가 없거나 임상 진료에 종사하지 않는 경우에는 참여 대상이 아닙니다.

3. 연구 방법: 참여에 동의하시면, 임상 5 단계(진료 전·검사 전·검사 후·진료 직후·진료 외 시간)에서 선호하는 AI 질의 유형(1·2 순위), 유형별 선호 답변 형식, AI 답변에 필요한 환자정보, AI 학습에서 제외해야 할 기록 유형, 연구·근거 질의 및 용어·약물 참조 질의의 사용 상황 등에 관한 익명 온라인 설문에 응답하시게 됩니다.

4. 연구 참여 기간 및 조사 시간: 1 회 응답이며, 소요 시간은 약 20~30 분입니다. (조사기간: 기관생명윤리위원회 승인일로부터 1 개월 이내)

5. 연구 참여 도중 중도탈락: 귀하는 설문을 제출하기 전까지 언제든지 응답을 중단할 수 있으며, 이로 인한 어떠한 불이익도 없습니다. 중단하신 경우 미완료 응답은 분석에서 제외됩니다. 다만 설문 제출 후에는 응답이 개인식별정보와 연결되지 않는 익명 자료로 저장되므로, 귀하의 응답을 특정하여 철회하거나 삭제하는 것은 어렵습니다.

6. 부작용 또는 위험요소: 본 설문은 비침습적 익명 조사로, 일상생활에서 통상적으로 경험하는 수준을 초과하는 위험은 예상되지 않으며 연구 참여에 따른 위험은 최소수준입니다.

7. 연구 참여에 따른 이익: 귀하에게 직접적인 의학적 이득은 없습니다. 다만 설문을 완료하신 분께는 연구 참여에 대한 감사의 표시로 3 만원 상당의 모바일 기프티콘을 드립니다. 사례품 수령을 원하시는 경우 설문 마지막 화면의 별도 링크에서 휴대전화번호를 남겨주시면 되며, 이 정보는 귀하의 설문 응답과 연결되지 않고 발송 완료 후 즉시 파기됩니다. 사례품 수령을 신청하지 않으셔도 어떠한 불이익도 없으며, 응답은 동일하게 분석에 포함됩니다. 귀하가 제공하는 응답은 수의 AI 시스템 설계에 대한 이해를 증진하는 데 기여합니다.

8. 연구에 참여하지 않을 시 불이익: 귀하는 본 연구에 참여하지 않을 자유가 있으며, 참여하지 않아도 어떠한 불이익도 없습니다.

9. 개인정보와 비밀보장: 본 설문은 성명·연락처·IP 등 개인식별정보를 수집하지 않는 익명 조사이며, 「개인정보 보호법」 제 23 조에 따른 민감정보를 수집하지 않습니다. 사례품 지급을 위한 연락처는 설문 응답과 분리된 별도 경로로 수집되며, 응답 내용과 어떠한 방식으로도 연결되지 않고 발송 완료 즉시 파기됩니다. 수집된 응답은 통계적으로만 처리되며, 학술지·학회 공개 시 개인을 식별할 수 있는 정보는 사용되지 않습니다. 국립창원대학교 기관생명윤리위원회는 관련 규정 범위 내에서 연구 자료를 열람할 수 있습니다. 연구 자료는 연구 종료 후 3 년간 보관한 뒤 폐기됩니다.

10. 이해관계 고지: 본 연구의 예비 문항은 ㈜춘옥컴퍼니가 운영한 시범 챗봇의 비식별 질의 로그를 검토하여 도출되었으며, 연구진은 동사 소속입니다. 본 연구에 대한 별도의 연구비 지원은 없으며, 사례품 지급 비용에 한하여 동사가 수행하는 중소벤처기업부 팁스(TIPS) 지원사업 재원에서 집행됩니다. 관련 이해상충은 국립창원대학교 기관생명윤리위원회에 신고되어 있습니다.

11. 연구 문의: 연구책임자 허혜연 / 이메일

※ 1 인 1 회 응답을 부탁드립니다. ※ 본 연구는 국립창원대학교 기관생명윤리위원회의 심의를 거쳐 수행됩니다. (승인번호: 7001066-202607-HR-053) ※ 본 설문에 응답·제출하시는 것은 본 연구 참여에 동의하시는 것으로 간주됩니다. 참여를 원하지 않으시면 이 창을 닫아 주십시오. 바쁘신 가운데 참여해 주셔서 감사합니다.

### 참여 자격 확인

**S1. 귀하는 수의사 면허를 보유하고 계십니까?**

☐ 예
☐ 아니오

**S2. 현재 임상 진료에 종사하고 계십니까?**

☐ 예
☐ 아니오

*두 문항 중 하나라도 "아니오"를 선택하신 경우, 본 연구의 참여 대상에 해당하지 않아 설문이 종료됩니다. ㈜춘옥컴퍼니 임직원인 경우에도 참여할 수 없습니다.*

### 배경 정보

**1. 소속 유형**

☐ 1 차 병원
☐ 2 차 병원
☐ 대학병원·교육기관
☐ 연구기관
☐ 기타(________)

**2. 임상 경력: 약 ____년 ____개월** *[서술형]*

**3. 주 진료분야(복수 기입 가능): ________** *[서술형]*

*질의유형 보기(Legend) — 임상(A–H): A 사실확인 · B 감별·추론 · C 치료·프로토콜 · D 임상 의사결정 지원 · E 차트·문서작성 · F 보호자 커뮤니케이션 · G 학술·근거 탐색 · H 교육·설명 / 연구(I–L): I 예후/치료효과 · J 약물/처치 근거 · K 감별/검사 해석 · L 문헌비교 / 용어·약물(M–U): M 현장 즉답 · N 임상 접근 · O 감별진단 · P 프로토콜 · Q 검사해석 · R 용량/투약 · S 영상해석 · T 시술 · U 질환/진단*

##### 모듈 1. 임상 질의 ㅡ 단계별 질의유형 및 선호 답변구조

**Q1. 진료 전 — AI 질문 유형 (1 순위 ____ 2 순위 ____ — 보기 번호 기입)** *[순위]*

☐ 1) 빠른 사실 확인(A 형)
☐ 2) 감별·추론(B 형)
☐ 3) 치료·프로토콜(C 형)
☐ 4) 임상 의사결정 지원(D 형)
☐ 5) 차트·문서 작성(E 형)
☐ 6) 보호자 커뮤니케이션(F 형)
☐ 7) 학술·근거 탐색(G 형)
☐ 8) 교육·설명 요청(H 형)
☐ 9) 기타(직접 기술)

**Q2. 진료 중(검사 전) — AI 질문 유형 (1 순위 ____ 2 순위 ____ — 보기 번호 기입)** *[순위]*

☐ 1) 빠른 사실 확인(A 형)
☐ 2) 감별·추론(B 형)
☐ 3) 치료·프로토콜(C 형)
☐ 4) 임상 의사결정 지원(D 형)
☐ 5) 차트·문서 작성(E 형)
☐ 6) 보호자 커뮤니케이션(F 형)
☐ 7) 학술·근거 탐색(G 형)
☐ 8) 교육·설명 요청(H 형)

**Q3. 진료 중(검사 후) — AI 질문 유형 (1 순위 ____ 2 순위 ____ — 보기 번호 기입)** *[순위]*

☐ 1) 빠른 사실 확인(A 형)
☐ 2) 감별·추론(B 형)
☐ 3) 치료·프로토콜(C 형)
☐ 4) 임상 의사결정 지원(D 형)
☐ 5) 차트·문서 작성(E 형)
☐ 6) 보호자 커뮤니케이션(F 형)
☐ 7) 학술·근거 탐색(G 형)
☐ 8) 교육·설명 요청(H 형)

**Q4. 진료 직후 — AI 질문 유형 (1 순위 ____ 2 순위 ____ — 보기 번호 기입)** *[순위]*

☐ 1) 빠른 사실 확인(A 형)
☐ 2) 감별·추론(B 형)
☐ 3) 치료·프로토콜(C 형)
☐ 4) 임상 의사결정 지원(D 형)
☐ 5) 차트·문서 작성(E 형)
☐ 6) 보호자 커뮤니케이션(F 형)
☐ 7) 학술·근거 탐색(G 형)
☐ 8) 교육·설명 요청(H 형)
☐ 9) 기타(직접 기술)

**Q5. 진료 외 시간 — AI 질문 유형 (1 순위 ____ 2 순위 ____ — 보기 번호 기입)** *[순위]*

☐ 1) 빠른 사실 확인(A 형)
☐ 2) 감별·추론(B 형)
☐ 3) 치료·프로토콜(C 형)
☐ 4) 임상 의사결정 지원(D 형)
☐ 5) 차트·문서 작성(E 형)
☐ 6) 보호자 커뮤니케이션(F 형)
☐ 7) 학술·근거 탐색(G 형)
☐ 8) 교육·설명 요청(H 형)

**Q6. 실제 임상 질문 예시** *[서술형]*

**Q7. A 형(사실 확인) 답변 구조 (1 개 선택)**

☐ ① 핵심 수치만 간결하게
☐ ② 수치 + 간단한 임상 맥락
☐ ③ 수치 + 정상범위 비교 + 해석
☐ ④ 기타

**Q8. B 형(감별·추론) 답변 구조 (1 개 선택)**

☐ ① DDx 리스트 + 각각 근거
☐ ② 요약 → 근거 → 수치 → 종합 → Next step
☐ ③ 우선순위 DDx + 확진 검사
☐ ④ 기타

**Q9. C 형(치료·프로토콜) 답변 구조 (1 개 선택)**

☐ ① 단계별 프로토콜
☐ ② 약물명 + 용량 + 주의사항 표
☐ ③ 프로토콜 + 모니터링 포인트

**Q10. D 형(의사결정 지원) 답변 구조 (1 개 선택)**

☐ ① 장단점 비교표
☐ ② 결론 먼저 + 근거 뒤에

**Q11. E 형(차트 작성) 답변 구조 (1 개 선택)**

☐ ① S/O/A/P 기본 구조만
☐ ② S/O/A/P + 영어·한국어 병기 + Plan 3 분류
☐ ③ S/O/A/P + 판단 근거 주석
☐ ④ 기타

**Q12. F 형(보호자 안내) 답변 구조 (1 개 선택)**

☐ ① 짧은 메시지형 — 2~3 줄, 핵심만 (카톡/문자)
☐ ② 중간 설명형 — 5~6 줄, 핵심 + 이유 + 주의사항
☐ ③ 상세 안내형 — 주의사항·일정·약 복용법까지 포함

**Q13. G 형(근거 탐색) 답변 구조 (1 개 선택)**

☐ ① 결론 + DOI 링크만
☐ ② 결론 + 출처 명시(논문명, 저자, 연도) + DOI 링크
☐ ③ 결론 + 출처 명시 + 근거 수준(연구 설계, 케이스 수, 신뢰도) + DOI 링크

**Q14. H 형(교육·설명) 답변 구조 (1 개 선택)**

☐ ① 핵심 기전 3~5 줄
☐ ② 기전 + 임상 연결
☐ ③ 교과서 수준 상세

**Q15. AI 답변 필수 포함 요소** *[서술형]*

**모듈 2. 임상 질의 — 필수 환자정보**

**Q1. AI 답변에 반드시 포함되어야 할 환자 정보 (해당하는 항목 모두 선택)** *[복수선택]*

☐ ① 종(개/고양이)
☐ ② 품종
☐ ③ 나이
☐ ④ 체중
☐ ⑤ 기저질환(기존 진단명)
☐ ⑥ 현재 복용 중인 약물
☐ ⑦ 중성화 여부
☐ ⑧ 보호자 경제 상황/치료 의지
☐ ⑨ 기타(직접 기술)

**Q2. 있으면 좋은 환자 정보 (우선순위대로 순위 기입: 1, 2, 3 …)** *[순위]*

☐ ① 종(개/고양이)
☐ ② 품종
☐ ③ 나이
☐ ④ 체중
☐ ⑤ 기저질환(기존 진단명)
☐ ⑥ 현재 복용 중인 약물
☐ ⑦ 중성화 여부
☐ ⑧ 보호자 경제 상황/치료 의지
☐ ⑨ 기타(직접 기술)

**Q3. 맥락 정보가 판단을 바꾼 경험** *[서술형]*

**모듈 3. 임상 질의 — 학습 제외 기록 및 편향 방지**

**Q1. AI 학습에서 제외해야 할 진료기록 유형 (해당하는 항목 모두 선택)** *[복수선택]*

☐ ① 보호자 비용 문제로 검사 거부 → 진단 없이 대증치료만 한 기록
☐ ② 응급이라 기록이 부실하거나 빠진 항목이 많은 기록
☐ ③ 오래된 프로토콜로 치료한 기록(현재 가이드라인과 다름)
☐ ④ 수련의/인턴이 작성하고 어텐딩 확인이 안 된 기록
☐ ⑤ 보호자 요청으로 표준이 아닌 치료를 한 기록
☐ ⑥ 기타(직접 기술)

**Q2. 데이터 편향 방지 방법 (해당하는 항목 모두 선택)** *[복수선택]*

☐ ① 다양한 지역(서울/지방)의 병원 데이터를 균등하게 확보
☐ ② 1 차 병원과 2·3 차 병원 데이터를 균형 있게 수집
☐ ③ 인기 품종 외에 드문 품종 데이터를 의도적으로 보강
☐ ④ 내과 외에 외과·안과·피부과 등 전문과목 데이터 확보
☐ ⑤ 해외 논문/가이드라인 데이터로 국내 데이터 부족분 보완
☐ ⑥ 기타(직접 기술)

**모듈 4. 연구·근거 질의 — 사용 상황 및 답변 우선순위**

**Q1. I 형(예후/치료효과) 사용 상황 (1 순위 ____ 2 순위 ____)** *[순위]*

☐ ① 보호자가 "얼마나 살 수 있나요?" 물어봤을 때
☐ ② 치료 계획을 세우면서 성공률/생존기간 비교할 때
☐ ③ 환자 상태 악화로 예후를 재평가해야 할 때
☐ ④ 수술 후 재발률이나 장기 예후를 확인할 때

**Q2. J 형(약물/처치) 사용 상황 (1 순위 ____ 2 순위 ____)** *[순위]*

☐ ① 처음 쓰는 약물의 용량·투여법을 확인할 때
☐ ② 기존 약물의 최신 권고 용량이 바뀌었는지 확인할 때
☐ ③ 부작용이 나타나서 대안 약물을 찾을 때
☐ ④ 두 약물 병용 시 상호작용이 궁금할 때
☐ ⑤ 기타(직접 기술)

**Q3. K 형(감별/검사) 사용 상황 (1 순위 ____ 2 순위 ____)** *[순위]*

☐ ① 검사 수치가 애매해서 해석이 필요할 때
☐ ② 증상이 비전형적이라 DDx 를 넓혀야 할 때
☐ ③ 영상 소견이 여러 진단을 가리킬 때
☐ ④ 드문 질환이 의심되어 감별 기준을 확인할 때
☐ ⑤ 기타(직접 기술)

**Q4. L 형(문헌비교) 사용 상황 (1 순위 ____ 2 순위 ____)** *[순위]*

☐ ① 치료법 A vs B 중 어떤 게 나은지 근거를 확인할 때
☐ ② 보호자가 인터넷에서 본 치료법의 근거를 확인할 때
☐ ③ 학회/세미나에서 들은 접근법의 근거를 찾을 때
☐ ④ 기존 프로토콜을 바꿔야 할지 판단 근거가 필요할 때
☐ ⑤ 기타(직접 기술)

**Q5. I~L 형 외 추가 질문 유형** *[서술형]*

**Q6. 논문 답변 시 정보 우선순위 (순위 기입)** *[순위]*

☐ ① 논문 기초 정보(저자, 저널명, 연구 연도 등)
☐ ② 결론/핵심 수치(생존기간, 성공률 등)
☐ ③ 대상군 정보(종, 품종, 마릿수 등)
☐ ④ 임상 적용 가능성(국내 환경에서 쓸 수 있는지)

**Q7. 근거 수준 표현 방식 (1 개 선택)**

☐ A — 출처를 각각 명시(논문명, 저자, 연도)
☐ B — 연구 규모를 명시(RCT 1 건, 138 마리 코호트 등)
☐ C — 신뢰도를 등급으로 표시(근거 강함/보통/약함)
☐ D — 기타(직접 작성)

**Q8. 근거 수준 위치 (1 개 선택)**

☐ A — 답변 앞
☐ B — 답변 뒤
☐ C — 문장 안에 자연스럽게
☐ D — 기타(직접 작성)

**Q9. 연구 근거 답변 구조 (1 개 선택)**

☐ A — 결론 먼저형
☐ B — PICO 순서형
☐ C — 비교표형
☐ D — 기타(직접 작성)

**Q10. 논문 인용 시 금지 사항** *[서술형]*

**Q11. 논문 답변 필수 포함 요소 (해당하는 항목 모두 선택)** *[복수선택]*

☐ ① 출처(논문명, 저자, 연도)
☐ ② 케이스 수(n 수)
☐ ③ 연구 대상 종/품종
☐ ④ 연구의 한계점
☐ ⑤ 국내 적용 가능 여부

#### 모듈 5. 연구·근거 질의 — 학습 포인트 및 답변 순서

**Q1. 임상 연구 — 학습 포인트 (순위 기입)** *[순위]*

☐ ① 연구 목적 + 방법 요약
☐ ② 대상군 정보(종, 품종, n 수)
☐ ③ 결과 수치(P-value, 생존기간 등)
☐ ④ 결론

**Q2. 약물/처치 연구 — 학습 포인트 (순위 기입)** *[순위]*

☐ ① 약물명(INN) + 용량 + 투여법
☐ ② 주요 부작용 + 모니터링 항목
☐ ③ 대상군 정보(종, 품종, n 수)
☐ ④ 결론 + 임상 권고사항

**Q3. 증례 보고 — 학습 포인트 (순위 기입)** *[순위]*

☐ ① 환자 시그널먼트(종, 품종, 나이)
☐ ② 주요 임상 소견 + 경과
☐ ③ 치료 방법 + 결과
☐ ④ 케이스 수 + 일반화 한계

**Q4. 기초/개념 연구 — 학습 포인트 (순위 기입)** *[순위]*

☐ ① 핵심 기전/경로 설명 부분
☐ ② 임상 적용 가능성 부분
☐ ③ 결론 + 향후 연구 방향
☐ ④ 참고한 주요 선행 연구 목록

**Q5. I 형 답변 순서 (순위 기입)** *[순위]*

☐ ① 생존기간/성공률 수치
☐ ② P-value 또는 통계적 유의성
☐ ③ 비교군 간 차이(치료군 vs 대조군)
☐ ④ 케이스 수(n) + 연구 설계
☐ ⑤ 임상 적용 한계(단일 증례 등)
☐ ⑥ 기타(직접 기술)

**Q6. J 형 답변 순서 (순위 기입)** *[순위]*

☐ ① 약물명(INN) + 구체적 용량
☐ ② 투여 경로 + 투여 간격
☐ ③ 주요 부작용 목록
☐ ④ 모니터링 항목(언제 무엇을 체크)
☐ ⑤ 금기/주의사항

**Q7. K 형 답변 순서 (순위 기입)** *[순위]*

☐ ① 감별진단 목록(가능성 순)
☐ ② 각 진단별 특징/구분점
☐ ③ 확진 방법(추천 검사)
☐ ④ 영문 용어 유지(ddx, head tilt 등)
☐ ⑤ 긴급도/우선 배제 대상 표시

**Q8. L 형 답변 순서 (순위 기입)** *[순위]*

☐ ① 각 선택지의 장단점 비교
☐ ② 비교표 형태(A vs B 나란히)
☐ ③ 근거가 되는 연구 출처
☐ ④ 근거 수준(얼마나 믿을 만한 연구인지) 비교
☐ ⑤ 임상 상황별 권고
☐ ⑥ 기타(직접 기술)

**Q9. 요약 길이/형태 (1 개 선택)**

☐ A — 1~2 문장 핵심만
☐ B — 3~5 문장 결론+수치+제한점
☐ C — 섹션별 구분
☐ D — 기타(직접 작성)

**Q10. 요약 필수 포함 요소** *[서술형]*

**Q11. 연구 근거 질의 관련 추가 의견** *[서술형]*

## 모듈 6. 연구·근거 질의 — 학습 제외 연구

**Q1. AI 학습에서 제외해야 할 연구 유형** *[서술형]*

**Q2. 해외 가이드라인을 국내에 적용하면서 어려움을 겪은 경험이 있다면 기술해 주십시오.** *[서술형]*

## 모듈 7. 용어·약물 참조 질의 — 사용 상황 및 답변 순서

**Q1. M 형(현장 즉답) 사용 상황 (1 순위 ____ 2 순위 ____)** *[순위]*

☐ ① 진료 중 처음 보는 용어가 나왔을 때
☐ ② 논문이나 교재를 읽다가 빠르게 확인할 때
☐ ③ 후배/실습생에게 설명해줘야 할 때
☐ ④ 기타(직접 기술)

**Q2. N 형(임상 접근) 사용 상황 (1 순위 ____ 2 순위 ____)** *[순위]*

☐ ① 익숙하지 않은 질환/상태의 환자가 왔을 때
☐ ② work-up 순서가 헷갈려서 확인할 때
☐ ③ 응급 상황에서 초기 접근법을 빠르게 확인할 때
☐ ④ 다른 과 영역(안과, 정형 등)의 접근법이 필요할 때

**Q3. O 형(감별진단) 사용 상황 (1 순위 ____ 2 순위 ____)** *[순위]*

☐ ① 놓치는 감별이 없는지 확인할 때
☐ ② 검사 결과가 여러 진단을 가리킬 때
☐ ③ 보호자에게 가능한 진단 목록을 설명해야 할 때
☐ ④ 흔하지 않은 감별까지 넓게 보고 싶을 때

**Q4. P 형(프로토콜) 사용 상황 (1 순위 ____ 2 순위 ____)** *[순위]*

☐ ① 처방 직전에 정확한 용량/투여 간격을 확인할 때
☐ ② 복합 처치의 순서나 타이밍을 정리할 때
☐ ③ 특수 환자(신부전, 간질환 등)에서 용량 조절이 필요할 때
☐ ④ 입원 환자의 모니터링 프로토콜을 세울 때
☐ ⑤ 기타(직접 기술)

**Q5. Q 형(검사 해석) 사용 상황 (1 순위 ____ 2 순위 ____)** *[순위]*

☐ ① 검사 결과지에서 익숙하지 않은 항목을 확인할 때
☐ ② 수치가 애매해서 정상/비정상 판단이 어려울 때
☐ ③ 여러 수치를 종합해서 해석이 필요할 때
☐ ④ 기타(직접 기술)

**Q6. R 형(용량/투약) 사용 상황 (1 순위 ____ 2 순위 ____)** *[순위]*

☐ ① 처음 쓰는 약물의 용량을 확인할 때
☐ ② 투여 간격(q8h, q12h 등)이 헷갈릴 때
☐ ③ CRI 등 특수 투여법의 계산이 필요할 때
☐ ④ 기타(직접 기술)

**Q7. S 형(영상 해석) 사용 상황 (1 순위 ____ 2 순위 ____)** *[순위]*

☐ ① 영상 소견을 해석해야 할 때
☐ ② 영상 검사 선택이 고민될 때(X-ray vs 초음파 vs CT 등)
☐ ③ 영상에서 정상 vs 비정상 구분이 필요할 때
☐ ④ 기타(직접 기술)

**Q8. T 형(시술) 사용 상황 (1 순위 ____ 2 순위 ____)** *[순위]*

☐ ① 시술 순서나 방법이 헷갈릴 때
☐ ② 처음 하는 시술의 과정을 확인할 때
☐ ③ 시술 중 합병증 대처법을 확인할 때
☐ ④ 기타(직접 기술)

**Q9. U 형(질환/진단) 사용 상황 (1 순위 ____ 2 순위 ____)** *[순위]*

☐ ① 드문 질환이 의심될 때
☐ ② 진단 기준을 정확히 확인하고 싶을 때
☐ ③ 보호자에게 예후를 설명해야 할 때
☐ ④ 기타(직접 기술)

**Q10. M 형 답변 순서 (순위 기입)** *[순위]*

☐ ① 정의(무엇인지)
☐ ② 정상 범위/기준치
☐ ③ 임상 의미(왜 중요한지)
☐ ④ 주의사항

**Q11. N 형 답변 순서 (순위 기입)** *[순위]*

☐ ① 초기 접근 순서(work-up)
☐ ② 필수 검사 항목
☐ ③ 감별진단 목록
☐ ④ 응급 여부 판단 기준

**Q12. O 형 답변 순서 (순위 기입)** *[순위]*

☐ ① 감별진단 목록(가능성 순)
☐ ② 각 진단별 구분 포인트
☐ ③ 확진 검사
☐ ④ 긴급 배제 대상

**Q13. P 형 답변 순서 (순위 기입)** *[순위]*

☐ ① 약물명 + 용량
☐ ② 투여 경로·간격
☐ ③ 모니터링 포인트
☐ ④ 금기/주의사항

**Q14. Q 형 답변 순서 (순위 기입)** *[순위]*

☐ ① 정상 범위
☐ ② 비정상의 임상 의미
☐ ③ 관련 감별진단
☐ ④ 추가 검사 권고

**Q15. R 형 답변 순서 (순위 기입)** *[순위]*

☐ ① 용량(mg/kg)
☐ ② 투여 경로·간격
☐ ③ 용량 조절 기준(신부전 등)
☐ ④ 부작용/모니터링

**Q16. S 형 답변 순서 (순위 기입)** *[순위]*

☐ ① 소견 설명
☐ ② 감별진단
☐ ③ 정상 vs 비정상 구분
☐ ④ 추가 검사/추적 권고

**Q17. T 형 답변 순서 (순위 기입)** *[순위]*

☐ ① 시술 순서/방법
☐ ② 준비물/장비
☐ ③ 합병증 대처
☐ ④ 주의사항/금기

**Q18. U 형 답변 순서 (순위 기입)** *[순위]*

☐ ① 정의/진단 기준
☐ ② 감별진단
☐ ③ 예후/경과
☐ ④ 치료 옵션 개요
☐ ⑤ 기타(직접 기술)

**Q19. 용어집 전체 답변 스타일 (1 개 선택)**

☐ A — 산문형(문장으로 서술)
☐ B — 구조화형([정의][용량][주의] 라벨 정리)
☐ C — 기타(직접 작성)

**Q20. 추가 의견** *[서술형]*

**설문 종료 화면**

설문이 완료되었습니다. 참여해 주셔서 감사합니다. 연구 참여에 대한 감사의 표시로 3 만원 상당의 모바일 기프티콘을 드립니다. 수령을 원하시면 아래 링크(본 설문과 분리된 별도 폼)에서 휴대전화번호를 남겨주십시오. 입력하신 번호는 위 설문 응답과 연결되지 않으며, 기프티콘 발송 완료 후 즉시 파기됩니다. 신청하지 않으셔도 귀하의 응답은 동일하게 연구에 활용되며 어떠한 불이익도 없습니다.

### Supplementary Material S4. Descriptive Results of Survey Items Not Reported in the Main Text

*Counts and percentages use all 43 respondents as the denominator, except in Table S4-1, where the number of valid second-choice responses at each clinical stage is used. For ranked items other than Table S4-1, only first choices are shown. For multiple-selection items, percentages may sum to more than 100%. Free-text entries under "Other" are counted with the "Other" option. These are descriptive summaries of self-reported preferences; they were not included in inferential analyses and do not measure actual query frequency. Question wording is abbreviated from Supplementary Material S3.*

#### Module 1 (Clinical L1): second-choice query types by clinical stage

##### Supplementary Table S4-1. Second-Choice Clinical Support Query Types Across Five Clinical Stages

| Clinical stage | Valid n | A | B | C | D | E | F | G | H |
|---|---|---|---|---|---|---|---|---|---|
| Pre-consultation | 37 | 1 (2.7%) | 8 (21.6%) | 10 (27.0%) | 6 (16.2%) | 2 (5.4%) | 0 | 7 (18.9%) | 3 (8.1%) |
| Intra-consultation (pre-test) | 37 | 1 (2.7%) | 4 (10.8%) | 12 (32.4%) | 8 (21.6%) | 0 | 1 (2.7%) | 7 (18.9%) | 4 (10.8%) |
| Intra-consultation (post-test) | 38 | 0 | 4 (10.5%) | 16 (42.1%) | 5 (13.2%) | 0 | 5 (13.2%) | 5 (13.2%) | 3 (7.9%) |
| Post-consultation | 37 | 0 | 1 (2.7%) | 7 (18.9%) | 3 (8.1%) | 2 (5.4%) | 9 (24.3%) | 13 (35.1%) | 2 (5.4%) |
| Off-duty | 39 | 1 (2.6%) | 3 (7.7%) | 8 (20.5%) | 5 (12.8%) | 1 (2.6%) | 0 | 13 (33.3%) | 8 (20.5%) |

Types: A, Quick fact check; B, Differential reasoning; C, Treatment/protocol; D, Clinical decision support; E, Chart/document writing; F, Client communication; G, Literature/evidence search; H, Education/explanation request. Percentages were calculated using the number of valid second-choice responses at each clinical stage as the denominator. Valid second-choice responses were available for 37, 37, 38, 37, and 39 respondents across the five stages, respectively; the corresponding numbers of respondents who did not record a second choice were 6, 6, 5, 6, and 4. First-choice responses for the same items are reported in Table 3 of the main text, where the stage denominators are 42, 43, 43, 42, and 43. Because the two sets of denominators differ, first- and second-choice counts should not be added together. Descriptively, Type C (Treatment/protocol) was the most frequent second choice from the pre-consultation stage through the intra-consultation (post-test) stage, whereas Type G (Literature/evidence search) was the most frequent second choice at the post-consultation and off-duty stages. These second-choice distributions were not included in any inferential analysis.

#### Module 1 (Clinical L1): answer structure for Types E, F, and H

##### M1 Q11. Answer structure for Type E (chart writing) (choose one) *(single choice)*

| Option | n | % |
|---|---|---|
| ① Basic S/O/A/P structure only | 15 | 34.9 |
| ② S/O/A/P + English–Korean terms + three-part Plan | 14 | 32.6 |
| ③ S/O/A/P + annotations explaining the reasoning | 13 | 30.2 |
| ④ Other | 1 | 2.3 |

##### M1 Q12. Answer structure for Type F (client guidance) (choose one) *(single choice)*

| Option | n | % |
|---|---|---|
| ① Short message — 2–3 lines, key points only (messenger/text) | 5 | 11.6 |
| ② Medium explanation — 5–6 lines, key point + reason + precautions | 21 | 48.8 |
| ③ Detailed guidance — including precautions, schedule, and how to give medication | 17 | 39.5 |

##### M1 Q14. Answer structure for Type H (education/explanation) (choose one) *(single choice)*

| Option | n | % |
|---|---|---|
| ① Core mechanism in 3–5 lines | 8 | 18.6 |
| ② Mechanism + clinical link | 29 | 67.4 |
| ③ Textbook-level detail | 6 | 14.0 |

#### Module 2 (Clinical L2)

##### M2 Q2. Patient information that would be good to have (rank in order of priority: 1, 2, 3 …) *(first choice)*

| Option | n | % |
|---|---|---|
| ① Species (dog/cat) | 4 | 9.3 |
| ② Breed | 11 | 25.6 |
| ③ Age | 8 | 18.6 |
| ④ Body weight | 2 | 4.7 |
| ⑤ Underlying disease (existing diagnoses) | 5 | 11.6 |
| ⑥ Current medications | 3 | 7.0 |
| ⑦ Neuter status | 2 | 4.7 |
| ⑧ Owner's financial situation/willingness to treat | 7 | 16.3 |
| ⑨ Other (please specify) | 1 | 2.3 |

#### Module 4 (Research L1): Types I–L

##### M4 Q1. Situations in which you would use Type I (prognosis/treatment effect) (1st ____ 2nd ____) *(first choice)*

| Option | n | % |
|---|---|---|
| ① When an owner asks, “How long will my pet live?” | 4 | 9.3 |
| ② When comparing success rates/survival times while planning treatment | 33 | 76.7 |
| ③ When reassessing prognosis because the patient has deteriorated | 4 | 9.3 |
| ④ When checking recurrence rates or long-term prognosis after surgery | 2 | 4.7 |

##### M4 Q2. Situations in which you would use Type J (drug/procedure) (1st ____ 2nd ____) *(first choice)*

| Option | n | % |
|---|---|---|
| ① When checking the dose and administration of a drug used for the first time | 34 | 79.1 |
| ② When checking whether the recommended dose of a familiar drug has changed | 6 | 14.0 |
| ③ When looking for an alternative drug because of adverse effects | 0 | 0.0 |
| ④ When wondering about interactions between two drugs used together | 1 | 2.3 |
| ⑤ Other (please specify) | 2 | 4.7 |

##### M4 Q3. Situations in which you would use Type K (differential/test) (1st ____ 2nd ____) *(first choice)*

| Option | n | % |
|---|---|---|
| ① When test values are equivocal and need interpretation | 10 | 23.3 |
| ② When atypical signs require broadening the DDx | 17 | 39.5 |

| Option | n | % |
|---|---|---|
| ③ When imaging findings point to several diagnoses | 6 | 14.0 |
| ④ When a rare disease is suspected and differential criteria need checking | 10 | 23.3 |
| ⑤ Other (please specify) | 0 | 0.0 |

##### M4 Q4. Situations in which you would use Type L (literature comparison) (1st ____ 2nd ____)

*(first choice)*

| Option | n | % |
|---|---|---|
| ① When checking the evidence on whether treatment A or B is better | 31 | 72.1 |
| ② When checking the evidence for a treatment the owner saw on the internet | 1 | 2.3 |
| ③ When looking for evidence on an approach heard at a conference/seminar | 2 | 4.7 |
| ④ When evidence is needed to decide whether to change an existing protocol | 8 | 18.6 |
| ⑤ Other (please specify) | 1 | 2.3 |

##### M4 Q6. Priority of information in answers citing papers (rank 1, 2, 3 …) *(first choice)*

| Option | n | % |
|---|---|---|
| ① Basic paper information (authors, journal, year) | 16 | 37.2 |
| ② Conclusion/key figures (survival time, success rate, etc.) | 22 | 51.2 |
| ③ Study population (species, breed, number of animals) | 0 | 0.0 |
| ④ Clinical applicability (usable in the domestic setting) | 5 | 11.6 |

##### M4 Q7. How should the level of evidence be expressed? (choose one) *(single choice)*

| Option | n | % |
|---|---|---|
| A — State each source (title, authors, year) | 13 | 30.2 |
| B — State the size of the study (e.g., 1 RCT, cohort of 138 animals) | 9 | 20.9 |
| C — Show reliability as a grade (strong/moderate/weak evidence) | 13 | 30.2 |
| D — Other (please write) | 8 | 18.6 |

##### M4 Q8. Where should the level of evidence appear? (choose one) *(single choice)*

| Option | n | % |
|---|---|---|
| A — Before the answer | 27 | 62.8 |
| B — After the answer | 10 | 23.3 |
| C — Naturally within the sentences | 5 | 11.6 |
| D — Other (please write) | 1 | 2.3 |

##### M4 Q9. Structure of answers based on research evidence (choose one) *(single choice)*

| Option | n | % |
|---|---|---|
| A — Conclusion first | 12 | 27.9 |
| B — PICO order | 2 | 4.7 |
| C — Comparison table | 28 | 65.1 |
| D — Other (please write) | 1 | 2.3 |

##### M4 Q11. Elements that must be included in answers citing papers (select all that apply)

*(selections (multiple))*

| Option | n | % |
|---|---|---|
| ① Source (title, authors, year) | 37 | 86.0 |
| ② Number of cases (n) | 38 | 88.4 |
| ③ Species/breed studied | 26 | 60.5 |
| ④ Limitations of the study | 29 | 67.4 |
| ⑤ Applicability in Korea | 18 | 41.9 |

#### Module 5 (Research L2)

##### M5 Q1. Clinical research — learning points (rank 1, 2, 3 …) *(first choice)*

| Option | n | % |
|---|---|---|
| ① Summary of aim + methods | 19 | 44.2 |
| ② Study population (species, breed, n) | 2 | 4.7 |
| ③ Result values (P-value, survival time, etc.) | 11 | 25.6 |
| ④ Conclusion | 11 | 25.6 |

##### M5 Q2. Drug/procedure research — learning points (rank) *(first choice)*

| Option | n | % |
|---|---|---|
| ① Drug name (INN) + dose + administration | 26 | 60.5 |
| ② Main adverse effects + monitoring items | 4 | 9.3 |
| ③ Study population (species, breed, n) | 3 | 7.0 |
| ④ Conclusion + clinical recommendations | 10 | 23.3 |

##### M5 Q3. Case reports — learning points (rank) *(first choice)*

| Option | n | % |
|---|---|---|
| ① Patient signalment (species, breed, age) | 11 | 25.6 |
| ② Main clinical findings + course | 17 | 39.5 |
| ③ Treatment + outcome | 10 | 23.3 |
| ④ Number of cases + limits to generalization | 5 | 11.6 |

##### M5 Q4. Basic/conceptual research — learning points (rank) *(first choice)*

| Option | n | % |
|---|---|---|
| ① Explanation of the key mechanism/pathway | 34 | 79.1 |
| ② Clinical applicability | 5 | 11.6 |
| ③ Conclusion + future research directions | 4 | 9.3 |
| ④ List of key prior studies cited | 0 | 0.0 |

##### M5 Q5. Answer order for Type I (rank) *(first choice)*

| Option | n | % |
|---|---|---|
| ① Survival time/success rate | 27 | 62.8 |
| ② P-value or statistical significance | 0 | 0.0 |
| ③ Difference between groups (treatment vs. control) | 9 | 20.9 |
| ④ Number of cases (n) + study design | 7 | 16.3 |

| Option | n | % |
|---|---|---|
| ⑤ Limits to clinical application (e.g., single case) | 0 | 0.0 |
| ⑥ Other (please specify) | 0 | 0.0 |

##### M5 Q6. Answer order for Type J (rank) *(first choice)*

| Option | n | % |
|---|---|---|
| ① Drug name (INN) + specific dose | 40 | 93.0 |
| ② Route + dosing interval | 1 | 2.3 |
| ③ List of main adverse effects | 1 | 2.3 |
| ④ Monitoring items (what to check and when) | 1 | 2.3 |
| ⑤ Contraindications/precautions | 0 | 0.0 |

##### M5 Q7. Answer order for Type K (rank) *(first choice)*

| Option | n | % |
|---|---|---|
| ① Differential diagnosis list (in order of likelihood) | 39 | 90.7 |
| ② Distinguishing features of each diagnosis | 1 | 2.3 |
| ③ How to confirm (recommended tests) | 0 | 0.0 |
| ④ Keep English terms (ddx, head tilt, etc.) | 0 | 0.0 |
| ⑤ Indicate urgency/diagnoses to rule out first | 3 | 7.0 |

##### M5 Q8. Answer order for Type L (rank) *(first choice)*

| Option | n | % |
|---|---|---|
| ① Pros and cons of each option | 13 | 30.2 |
| ② Comparison table (A vs. B side by side) | 23 | 53.5 |
| ③ Source studies for the evidence | 3 | 7.0 |
| ④ Comparison of the level of evidence (how reliable the studies are) | 2 | 4.7 |
| ⑤ Recommendations by clinical situation | 2 | 4.7 |
| ⑥ Other (please specify) | 0 | 0.0 |

##### M5 Q9. Length/form of summaries (choose one) *(single choice)*

| Option | n | % |
|---|---|---|
| A — Key point only, in 1–2 sentences | 1 | 2.3 |
| B — Conclusion + figures + limitations, in 3–5 sentences | 29 | 67.4 |
| C — Divided into sections | 7 | 16.3 |
| D — Other (please write) | 6 | 14.0 |

#### Module 7 (Terminology L1): Types M–U

##### M7 Q1. Situations in which you would use Type M (instant answer) (1st ____ 2nd ____) *(first choice)*

| Option | n | % |
|---|---|---|
| ① When an unfamiliar term comes up during a consultation | 19 | 44.2 |
| ② When checking quickly while reading a paper or textbook | 21 | 48.8 |
| ③ When explaining to a junior colleague/student | 1 | 2.3 |
| ④ Other (please specify) | 2 | 4.7 |

**M7 Q2. Type N (clinical approach) (1st ____ 2nd ____)** *(first choice)*

| Option | n | % |
|---|---|---|
| ① When a patient with an unfamiliar disease/condition presents | 25 | 58.1 |
| ② When checking the work-up order because it is unclear | 1 | 2.3 |
| ③ When quickly checking the initial approach in an emergency | 8 | 18.6 |
| ④ When an approach from another field (ophthalmology, orthopedics, etc.) is needed | 9 | 20.9 |

**M7 Q3. Type O (differential diagnosis) (1st ____ 2nd ____)** *(first choice)*

| Option | n | % |
|---|---|---|
| ① When checking that no differential has been missed | 22 | 51.2 |
| ② When test results point to several diagnoses | 15 | 34.9 |
| ③ When explaining the list of possible diagnoses to an owner | 1 | 2.3 |
| ④ When wanting to consider uncommon differentials broadly | 5 | 11.6 |

**M7 Q4. Type P (protocol) (1st ____ 2nd ____)** *(first choice)*

| Option | n | % |
|---|---|---|
| ① When checking the exact dose/interval just before prescribing | 22 | 51.2 |
| ② When organizing the order or timing of combined treatments | 6 | 14.0 |
| ③ When dose adjustment is needed in special patients (renal failure, liver disease, etc.) | 13 | 30.2 |
| ④ When setting up a monitoring protocol for inpatients | 0 | 0.0 |
| ⑤ Other (please specify) | 2 | 4.7 |

**M7 Q5. Type Q (test interpretation) (1st ____ 2nd ____)** *(first choice)*

| Option | n | % |
|---|---|---|
| ① When checking an unfamiliar item on a test report | 13 | 30.2 |
| ② When values are equivocal and normal/abnormal is hard to judge | 15 | 34.9 |
| ③ When several values must be interpreted together | 15 | 34.9 |
| ④ Other (please specify) | 0 | 0.0 |

**M7 Q6. Type R (dose/administration) (1st ____ 2nd ____)** *(first choice)*

| Option | n | % |
|---|---|---|
| ① When checking the dose of a drug used for the first time | 32 | 74.4 |
| ② When the dosing interval (q8h, q12h, etc.) is unclear | 3 | 7.0 |
| ③ When calculations for special administration such as CRI are needed | 6 | 14.0 |
| ④ Other (please specify) | 2 | 4.7 |

**M7 Q7. Type S (imaging interpretation) (1st ____ 2nd ____)** *(first choice)*

| Option | n | % |
|---|---|---|
| ① When imaging findings need to be interpreted | 23 | 53.5 |
| ② When deciding which imaging test to choose (X-ray vs. ultrasound vs. CT, etc.) | 6 | 14.0 |
| ③ When normal vs. abnormal must be distinguished on imaging | 11 | 25.6 |
| ④ Other (please specify) | 3 | 7.0 |

**M7 Q8. Type T (procedure) (1st ____ 2nd ____)** *(first choice)*

| Option | n | % |
|---|---|---|
| ① When the order or method of a procedure is unclear | 6 | 14.0 |
| ② When checking the steps of a procedure performed for the first time | 29 | 67.4 |
| ③ When checking how to manage complications during a procedure | 6 | 14.0 |
| ④ Other (please specify) | 2 | 4.7 |

**M7 Q9. Type U (disease/diagnosis) (1st ____ 2nd ____)** *(first choice)*

| Option | n | % |
|---|---|---|
| ① When a rare disease is suspected | 18 | 41.9 |
| ② When wanting to check diagnostic criteria exactly | 16 | 37.2 |
| ③ When prognosis must be explained to an owner | 8 | 18.6 |
| ④ Other (please specify) | 1 | 2.3 |

**M7 Q10. Answer order for Type M (rank)** *(first choice)*

| Option | n | % |
|---|---|---|
| ① Definition (what it is) | 39 | 90.7 |
| ② Normal range/reference value | 4 | 9.3 |
| ③ Clinical significance (why it matters) | 0 | 0.0 |
| ④ Precautions | 0 | 0.0 |

**M7 Q11. Answer order for Type N (rank)** *(first choice)*

| Option | n | % |
|---|---|---|
| ① Initial approach order (work-up) | 33 | 76.7 |
| ② Essential tests | 4 | 9.3 |
| ③ Differential diagnosis list | 3 | 7.0 |
| ④ Criteria for judging urgency | 3 | 7.0 |

**M7 Q12. Answer order for Type O (rank)** *(first choice)*

| Option | n | % |
|---|---|---|
| ① Differential diagnosis list (in order of likelihood) | 38 | 88.4 |
| ② Distinguishing points of each diagnosis | 1 | 2.3 |
| ③ Confirmatory tests | 1 | 2.3 |
| ④ Diagnoses to rule out urgently | 3 | 7.0 |

**M7 Q13. Answer order for Type P (rank)** *(first choice)*

| Option | n | % |
|---|---|---|
| ① Drug name + dose | 39 | 90.7 |
| ② Route and interval | 3 | 7.0 |
| ③ Monitoring points | 1 | 2.3 |
| ④ Contraindications/precautions | 0 | 0.0 |

##### M7 Q14. Answer order for Type Q (rank) *(first choice)*

| Option | n | % |
|---|---|---|
| ① Normal range | 18 | 41.9 |
| ② Clinical significance of abnormality | 16 | 37.2 |
| ③ Related differential diagnoses | 7 | 16.3 |
| ④ Recommended additional tests | 2 | 4.7 |

##### M7 Q15. Answer order for Type R (rank) *(first choice)*

| Option | n | % |
|---|---|---|
| ① Dose (mg/kg) | 41 | 95.3 |
| ② Route and interval | 1 | 2.3 |
| ③ Criteria for dose adjustment (renal failure, etc.) | 1 | 2.3 |
| ④ Adverse effects/monitoring | 0 | 0.0 |

##### M7 Q16. Answer order for Type S (rank) *(first choice)*

| Option | n | % |
|---|---|---|
| ① Description of findings | 36 | 83.7 |
| ② Differential diagnoses | 4 | 9.3 |
| ③ Normal vs. abnormal | 3 | 7.0 |
| ④ Additional tests/follow-up recommendations | 0 | 0.0 |

##### M7 Q17. Answer order for Type T (rank) *(first choice)*

| Option | n | % |
|---|---|---|
| ① Procedure order/method | 26 | 60.5 |
| ② Supplies/equipment | 16 | 37.2 |
| ③ Management of complications | 1 | 2.3 |
| ④ Precautions/contraindications | 0 | 0.0 |

##### M7 Q18. Answer order for Type U (rank) *(first choice)*

| Option | n | % |
|---|---|---|
| ① Definition/diagnostic criteria | 21 | 48.8 |
| ② Differential diagnoses | 0 | 0.0 |
| ③ Prognosis/course | 21 | 48.8 |
| ④ Overview of treatment options | 1 | 2.3 |
| ⑤ Other (please specify) | 0 | 0.0 |

##### M7 Q19. Overall answer style for terminology queries (choose one) *(single choice)*

| Option | n | % |
|---|---|---|
| A — Prose (written in sentences) | 13 | 30.2 |
| B — Structured ([Definition][Dose][Precautions] labels) | 24 | 55.8 |
| C — Other (please write) | 6 | 14.0 |

### Supplementary Material S5. GEE Analysis Code and Output

*Development of an Exploratory Taxonomy of Veterinary AI Query Types and Stage-Specific Query Preferences: An Expert Panel Study (Manuscript ID 1899253)*

*Analysis scope. Binary participant-clustered GEE models used a logit link, an exchangeable working correlation structure, and robust sandwich covariance. The main A-H analyses excluded the two submitted non-use responses coded OTHER. The institution comparison excluded the two research-institution participants. Odds ratios are reported with unadjusted 95% Wald confidence intervals; Benjamini-Hochberg adjustment was applied across the four reported coefficient contrasts.*

*Reproducibility. The analyses used statsmodels 0.15.0, and the code below is written for that environment. The survey response dataset is not embedded here: the IRB-approved consent procedure described statistical processing and a three-year retention period rather than public release, so the dataset is available from the corresponding author on reasonable request, as stated in the Data Availability Statement.*

#### S5.1 Expected long-format input

| Column | Documented value |
|---|---|
| participant_id | Anonymous participant identifier, repeated across five stage rows |
| institution | secondary_referral / university_teaching / primary_care / research_institution |
| experience_years | Numeric clinical experience in years |
| stage | 1-5: pre-consultation, intra-consultation pre-test, intra-consultation post-test, post-consultation, off-duty |
| first_choice | A-H, or OTHER for the submitted non-use response |

#### S5.2 Analysis code

```
# Supplementary Material S5 - GEE analysis code
# Target environment reported in the manuscript: statsmodels 0.15.0
# Input: de-identified long-format CSV with one row per participant x clinical stage.
# Required columns:
# participant_id, institution, experience_years, stage, first_choice
# institution values: secondary_referral, university_teaching, primary_care, research_institution
# stage values: 1=pre-consultation, 2=intra-consultation pre-test,
#               3=intra-consultation post-test, 4=post-consultation, 5=off-duty
# first_choice values: A-H or OTHER

import csv
import numpy as np
import statsmodels.api as sm
from statsmodels.genmod.cov_struct import Exchangeable, Independence
from statsmodels.stats.multitest import multipletests

INPUT_CSV = "survey_analysis_input.csv"


def load_rows(path):
    rows = []
    with open(path, newline="", encoding="utf-8-sig") as f:
        for d in csv.DictReader(f):
            rows.append({
                "participant_id": d["participant_id"],
                "institution": d["institution"],
                "experience_years": float(d["experience_years"]),
                "exp10": int(float(d["experience_years"]) >= 10.0),
                "stage": int(d["stage"]),
                "first_choice": d["first_choice"].strip().upper(),
            })
    return rows


def fit_gee(rows, outcome_code, predictors, cov_struct=None, cov_type="robust"):
    """predictors = list of (name, function(row)->0/1 or numeric)."""
    y = np.array([int(r["first_choice"] == outcome_code) for r in rows], dtype=float)
    X_cols = [np.ones(len(rows), dtype=float)]
    names = ["Intercept"]
    for name, fn in predictors:
        X_cols.append(np.array([fn(r) for r in rows], dtype=float))
        names.append(name)
    X = np.column_stack(X_cols)
    groups = np.array([r["participant_id"] for r in rows])
```

```
    if cov_struct is None:
        cov_struct = Exchangeable()
    model = sm.GEE(
        endog=y,
        exog=X,
        groups=groups,
        family=sm.families.Binomial(),
        cov_struct=cov_struct,
    )
    result = model.fit(cov_type=cov_type)
    return result, names


def effect_row(result, names, term):
    i = names.index(term)
    ci = result.conf_int()[i]
    return {
        "beta": result.params[i],
        "SE": result.bse[i],
        "OR": np.exp(result.params[i]),
        "CI_low": np.exp(ci[0]),
        "CI_high": np.exp(ci[1]),
        "p": result.pvalues[i],
    }


def show(label, effect):
    print(
        f"{label}: beta={effect['beta']:.6f}, SE={effect['SE']:.6f}, "
        f"OR={effect['OR']:.6f}, 95% CI={effect['CI_low']:.6f}-{effect['CI_high']:.6f}, "
        f"p={effect['p']:.6g}"
    )


rows = load_rows(INPUT_CSV)

# Main A-H analysis excludes submitted non-use responses coded OTHER.
main = [r for r in rows if r["first_choice"] != "OTHER"]
institution_sample = [r for r in main if r["institution"] != "research_institution"]

is_exp10 = ("exp10", lambda r: r["exp10"])
is_uni = ("university_teaching", lambda r: int(r["institution"] == "university_teaching"))
is_primary = ("primary_care", lambda r: int(r["institution"] == "primary_care"))
stage_terms = [
    ("stage_2", lambda r: int(r["stage"] == 2)),
    ("stage_3", lambda r: int(r["stage"] == 3)),
```

```
    ("stage_4", lambda r: int(r["stage"] == 4)),
    ("stage_5", lambda r: int(r["stage"] == 5)),
]

# ------------------------------------------------------------------
# Primary models
# ------------------------------------------------------------------
G_exp, G_exp_names = fit_gee(main, "G", [is_exp10])
D_exp, D_exp_names = fit_gee(main, "D", [is_exp10])
G_inst, G_inst_names = fit_gee(institution_sample, "G", [is_uni, is_primary])

print("PRIMARY MODELS")
show("Type G, >=10 vs <10 years", effect_row(G_exp, G_exp_names, "exp10"))
print(f"  exchangeable alpha={float(G_exp.cov_struct.dep_params):.6f}")
show("Type D, >=10 vs <10 years", effect_row(D_exp, D_exp_names, "exp10"))
print(f"  exchangeable alpha={float(D_exp.cov_struct.dep_params):.6f}")
show("Type G, university/teaching vs secondary referral", effect_row(G_inst, G_inst_names, "university_teaching"))
show("Type G, primary care vs secondary referral", effect_row(G_inst, G_inst_names, "primary_care"))
print(f"  exchangeable alpha={float(G_inst.cov_struct.dep_params):.6f}")

R = np.array([[0, 1, 0], [0, 0, 1]], dtype=float)
wald = G_inst.wald_test(R, scalar=True)
print(f"Institution overall Wald chi2(2)={float(wald.statistic):.6f}, p={float(wald.pvalue):.6g}")

# BH correction across the four reported coefficient contrasts, in this order:
# G by experience, D by experience, university vs secondary, primary vs secondary.
raw_p = np.array([
    G_exp.pvalues[G_exp_names.index("exp10")],
    D_exp.pvalues[D_exp_names.index("exp10")],
    G_inst.pvalues[G_inst_names.index("university_teaching")],
    G_inst.pvalues[G_inst_names.index("primary_care")],
])
BH = multipletests(raw_p, method="fdr_bh")[1]
print("BH-adjusted p-values:", [float(x) for x in BH])

# ------------------------------------------------------------------
# Sensitivity analyses
# ------------------------------------------------------------------
print("\nSENSITIVITY: independence working correlation")
for label, sample, outcome, pred in [
    ("G experience", main, "G", [is_exp10]),
    ("D experience", main, "D", [is_exp10]),
    ("G institution", institution_sample, "G", [is_uni, is_primary]),
]:
    res, names = fit_gee(sample, outcome, pred, cov_struct=Independence())
    if label == "G institution":
```

```
        show(label + " - university", effect_row(res, names, "university_teaching"))
        show(label + " - primary", effect_row(res, names, "primary_care"))
    else:
        show(label, effect_row(res, names, "exp10"))

print("\nSENSITIVITY: bias-reduced covariance")
for label, sample, outcome, pred in [
    ("G experience", main, "G", [is_exp10]),
    ("D experience", main, "D", [is_exp10]),
    ("G institution", institution_sample, "G", [is_uni, is_primary]),
]:
    res, names = fit_gee(sample, outcome, pred, cov_struct=Exchangeable(), cov_type="bias_reduced")
    if label == "G institution":
        show(label + " - university", effect_row(res, names, "university_teaching"))
        show(label + " - primary", effect_row(res, names, "primary_care"))
    else:
        show(label, effect_row(res, names, "exp10"))

print("\nSENSITIVITY: adjustment for clinical stage (stage 1 reference)")
for label, sample, outcome, pred, term in [
    ("G experience + stage", main, "G", [is_exp10] + stage_terms, "exp10"),
    ("D experience + stage", main, "D", [is_exp10] + stage_terms, "exp10"),
    ("G institution + stage, university", institution_sample, "G", [is_uni, is_primary] + stage_terms, "university_teaching"),
    ("G institution + stage, primary", institution_sample, "G", [is_uni, is_primary] + stage_terms, "primary_care"),
]:
    res, names = fit_gee(sample, outcome, pred)
    show(label, effect_row(res, names, term))

print("\nSENSITIVITY: submitted non-use responses coded as non-selection")
all_rows = rows
all_institution = [r for r in rows if r["institution"] != "research_institution"]
for label, sample, outcome, pred, term in [
    ("G experience", all_rows, "G", [is_exp10], "exp10"),
    ("D experience", all_rows, "D", [is_exp10], "exp10"),
    ("G institution, university", all_institution, "G", [is_uni, is_primary], "university_teaching"),
    ("G institution, primary", all_institution, "G", [is_uni, is_primary], "primary_care"),
]:
    res, names = fit_gee(sample, outcome, pred)
    show(label, effect_row(res, names, term))

print("\nSENSITIVITY: same-sample experience model (41 participants used in institution analysis)")
res, names = fit_gee(institution_sample, "G", [is_exp10])
show("G experience, 41-participant sample", effect_row(res, names, "exp10"))

print("\nSENSITIVITY: joint model (experience + institution + stage)")
res, names = fit_gee(institution_sample, "G", [is_exp10, is_uni, is_primary] + stage_terms)
```

```
for term in ["exp10", "university_teaching", "primary_care", "stage_2", "stage_3", "stage_4", "stage_5"]:
    show("Joint model - " + term, effect_row(res, names, term))
```

#### S5.3 Output from the 43-participant analysis dataset

```
PRIMARY MODELS
Type G, >=10 vs <10 years: beta=1.127418, SE=0.522336, OR=3.087675, 95% CI=1.109221-8.594983, p=0.0308956
  exchangeable alpha=0.137534
Type D, >=10 vs <10 years: beta=-0.337462, SE=0.398625, OR=0.713579, 95% CI=0.326687-1.558664, p=0.397238
  exchangeable alpha=0.077477
Type G, university/teaching vs secondary referral: beta=2.227169, SE=0.585542, OR=9.273576, 95% CI=2.943290-29.218739, p=0.000142607
Type G, primary care vs secondary referral: beta=-0.277632, SE=0.722385, OR=0.757576, 95% CI=0.183878-3.121200, p=0.700737
  exchangeable alpha=0.041227
Institution overall Wald chi2(2)=16.799440, p=0.00022493
BH-adjusted p-values: [0.06179120735836853, 0.5296502942343014, 0.0005704273235945619, 0.7007366027502882]

SENSITIVITY: independence working correlation
G experience: beta=1.109423, SE=0.522959, OR=3.032609, 95% CI=1.088110-8.452007, p=0.0338853
D experience: beta=-0.333189, SE=0.398420, OR=0.716634, 95% CI=0.328218-1.564708, p=0.402999
G institution - university: beta=2.220347, SE=0.587423, OR=9.210526, 95% CI=2.912519-29.127298, p=0.00015695
G institution - primary: beta=-0.277632, SE=0.722385, OR=0.757576, 95% CI=0.183878-3.121200, p=0.700737

SENSITIVITY: bias-reduced covariance
G experience: beta=1.127418, SE=0.548088, OR=3.087675, 95% CI=1.054625-9.039928, p=0.039686
D experience: beta=-0.337462, SE=0.418062, OR=0.713579, 95% CI=0.314476-1.619188, p=0.419548
G institution - university: beta=2.227169, SE=0.660852, OR=9.273576, 95% CI=2.539389-33.866098, p=0.000751273
G institution - primary: beta=-0.277632, SE=0.822393, OR=0.757576, 95% CI=0.151148-3.797070, p=0.735672

SENSITIVITY: adjustment for clinical stage (stage 1 reference)
G experience + stage: beta=1.258115, SE=0.582635, OR=3.518784, 95% CI=1.123190-11.023820, p=0.0308226
D experience + stage: beta=-0.577937, SE=0.502053, OR=0.561055, 95% CI=0.209728-1.500905, p=0.249671
G institution + stage, university: beta=2.703166, SE=0.725733, OR=14.926916, 95% CI=3.599357-61.903511, p=0.000195518
G institution + stage, primary: beta=-0.211524, SE=0.800260, OR=0.809350, 95% CI=0.168637-3.884357, p=0.791534

SENSITIVITY: submitted non-use responses coded as non-selection
G experience: beta=1.087917, SE=0.518996, OR=2.968085, 95% CI=1.073263-8.208171, p=0.0360648
D experience: beta=-0.354243, SE=0.399846, OR=0.701705, 95% CI=0.320483-1.536399, p=0.375646
G institution, university: beta=2.120264, SE=0.564366, OR=8.333333, 95% CI=2.756956-25.188816, p=0.000172034
G institution, primary: beta=-0.277632, SE=0.722385, OR=0.757576, 95% CI=0.183878-3.121200, p=0.700737

SENSITIVITY: same-sample experience model (41 participants used in institution analysis)
G experience, 41-participant sample: beta=1.008649, SE=0.537028, OR=2.741893, 95% CI=0.957044-7.855416, p=0.0603529

SENSITIVITY: joint model (experience + institution + stage)
Joint model - exp10: beta=0.551069, SE=0.578607, OR=1.735107, 95% CI=0.558232-5.393087, p=0.340891
Joint model - university_teaching: beta=2.492446, SE=0.751016, OR=12.090819, 95% CI=2.774532-52.689214, p=0.000904162
```

```
Joint model - primary_care: beta=-0.395965, SE=0.803044, OR=0.673030, 95% CI=0.139471-3.247781, p=0.621956
Joint model - stage_2: beta=1.073128, SE=0.794076, OR=2.924514, 95% CI=0.616787-13.866677, p=0.176562
Joint model - stage_3: beta=-0.650046, SE=0.876655, OR=0.522022, 95% CI=0.093644-2.910039, p=0.458386
Joint model - stage_4: beta=-1.287983, SE=0.853147, OR=0.275827, 95% CI=0.051813-1.468372, p=0.131124
Joint model - stage_5: beta=2.175949, SE=0.572881, OR=8.810539, 95% CI=2.866590-27.079423, p=0.000145717
```

#### S5.4 Primary estimates reported in the manuscript

| Contrast | OR | 95% CI | Raw p | BH-adjusted p | Sample |
|---|---|---|---|---|---|
| Type G: >=10 vs <10 years | 3.09 | 1.11-8.59 | 0.031 | 0.062 | 43 participants; 213 analyzable A-H stage responses |
| Type D: >=10 vs <10 years | 0.71 | 0.33-1.56 | 0.397 | 0.530 | 43 participants; 213 analyzable A-H stage responses |
| Type G: university/teaching vs secondary referral | 9.27 | 2.94-29.22 | <0.001 | <0.001 | 41 participants; research-institution participants excluded |
| Type G: primary care vs secondary referral | 0.76 | 0.18-3.12 | 0.701 | 0.701 | 41 participants; research-institution participants excluded |

The institution factor as a whole was evaluated with a two-degree-of-freedom Wald test: chi-square(2) = 16.80, $p < 0.001$.

### Supplementary Material S6. Schematic of the Deployed Chatbot Interface

*Development of an Exploratory Taxonomy of Veterinary AI Query Types and Stage-Specific Query Preferences: An Expert Panel Study (Manuscript ID 1899253)*

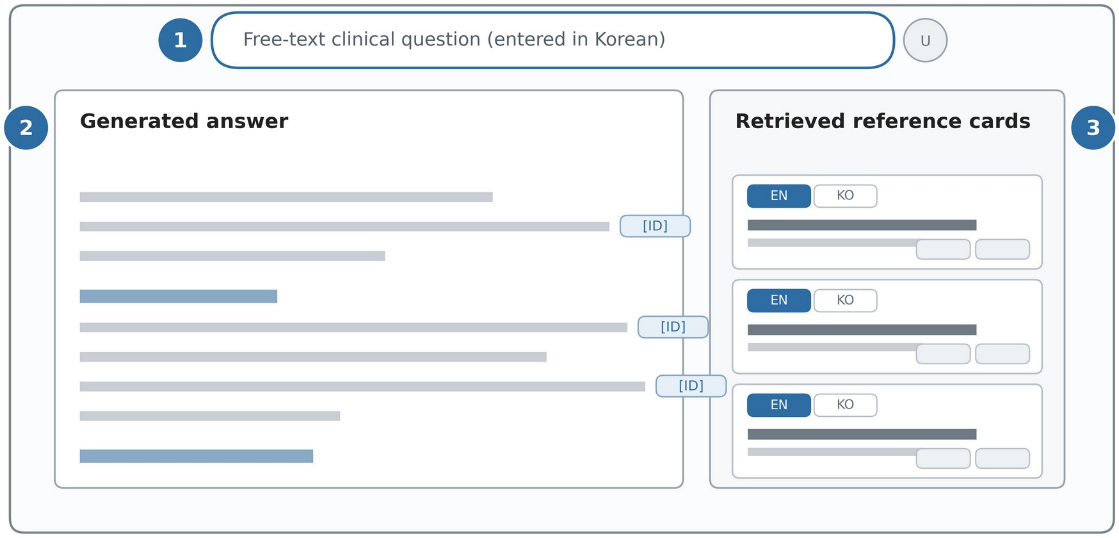


*Schematic only. Proportions and wording are illustrative; no logged query or user account is shown.*

*Figure S6. Layout of the retrieval-augmented chatbot as deployed at the three proof-of-concept sites during the log-collection period (1 August 2025 to 30 March 2026). (1) A single free-text box in which the veterinarian typed a clinical question in Korean. (2) The generated answer, presented as continuous Korean text under short headings, with inline markers linking statements to the retrieved sources. (3) A column of reference cards for the retrieved sources, each showing the article title and journal with a toggle between the English record and its Korean rendering. This is a schematic redrawn from an archived screenshot: that version of the interface is no longer in service, and the only surviving capture is too low in resolution to reproduce here. Proportions and labels are illustrative, and no logged query, account name, or product branding is shown. The interface was not an object of study; it is described here only so that readers can picture how the queries used in Phase 1 were produced.*
